# Consolidating species boundaries in the *Fusarium oxysporum* species complex based on genomic evidence

**DOI:** 10.64898/2026.09.25.754427

**Authors:** J. Bell, J. van Os, M. Thines, M.M. Costa, J. Collemare, J.Z. Groenewald, B. Mishra, B. Ulaszewski, P.W. Crous, M.F. Seidl

**Affiliations:** Theoretical Biology & Bioinformatics, Department of Biology, Utrecht University, Utrecht 3584 CH, The Netherlands; Senckenberg Biodiversity and Climate Research Center, Senckenberganlage 25, D-60325, Frankfurt am Main, Germany; Department of Biological Sciences, Institute of Ecology, Evolution and Diversity, Goethe-Universität Frankfurt am Main, Max-von-Laue Str. 13, D-60438, Frankfurt am Main, Germany; Westerdijk Fungal Biodiversity Institute, Uppsalalaan 8, 3584 CT, Utrecht, The Netherlands; Department of Genetics, Faculty of Biological Sciences, Kazimierz Wielki University in Bydgoszcz, Chodkiewicza 30, 85-064 Bydgoszcz, Poland; Microbiology, Department of Biology, Utrecht University, Padualaan 8, 3584 CH Utrecht The Netherlands

**Keywords:** Fusarium oxysporum, plant pathogens, phylogenomics, systematics, taxonomy

## Abstract

The *Fusarium oxysporum* species complex (FOSC) comprises numerous plant pathogens infecting crops such as tomato, lettuce, melon, and banana. Despite its economic and ecological importance and decades of research, the species names linked to different phylogenetic lineages within this species complex remain largely controversial. Earlier taxonomic frameworks relied on a limited number of marker genes, variable taxon sampling and inconsistent species definitions, leading to a fragmented and unstable taxonomy. To address these challenges, we here analysed nearly 2,500 conserved marker genes from 660 genomes, including genome assemblies from 22 ex-type strains that were sequenced using long-read sequencing technology, representing nearly half of the 51 currently described species within FOSC. Maximum likelihood- and coalescence-based analyses of the full phylogenomic data matrix as well as subsets yielded a robust phylogeny of FOSC, which was furthermore corroborated by whole genome comparisons. Many previously described species are redundant, while two novel lineages remain undescribed. Direct comparisons to traditional marker gene-based approaches demonstrate their inadequacy for species delimitation in FOSC. We show that our phylogenomic framework can guide the detection of novel marker genes able to distinguish the major lineages within FOSC, which will be essential for taxon delineation and classification in future studies. Thus, our phylogenomic analyses provide a stable cornerstone for revising FOSC taxonomy and offers a critical resource for the broader research community.

## INTRODUCTION

The *Fusarium oxysporum* species complex (FOSC) is one of the most intensively studied and economically important groups of filamentous fungi and belongs to the 4^th^ most cited genera of fungi (Bhunjun *et al*. 2024). Members of the FOSC display remarkable ecological versatility, encompassing saprotrophic, endophytic, and opportunistic human-associated lifestyles, but most notably include many phytopathogens (Edel-Hermann & Lecomte 2019, Martínez-Soto *et al*. 2023). Although the majority of FOSC isolates appear to be putatively non-pathogenic, research has traditionally focused on pathogenic lineages because of their agricultural and societal significance. Collectively, the phytopathogenic members of this species complex can infect more than 100 plant species, including key food and economic crops such as tomato, soybean, melon, cucumber, chickpea, lettuce, and banana (Edel-Hermann & Lecomte 2019), posing a serious and persistent threat to global food security. Consequently, *F. oxysporum* was identified as the fifth most important fungal plant pathogen worldwide (Dean *et al*. 2012).

Accurate species identification is crucial for diverse users, including farmers, ecologists, conservation officers, customs authorities, and phytopathologists. For instance, rapid and reliable identification of a pathogen underpins timely and effective disease management decisions (Hariharan & Prasannath 2021). Beyond its practical applications, a stable and consistent taxonomic framework also enables clear communication among scientists, provides a foundation for developing targeted strategies in disease control and ecological research, and is instrumental for evolutionary research (Crous *et al*. 2015, Jiménez-Mejías *et al*. 2024). The FOSC represents a phylogenetic clade of taxa that are morphologically indistinguishable, yet genetically distinct (Edel-Hermann & Lecomte 2019), making the delineation of species boundaries challenging. Such species complexes often arise through convergent evolution, recent divergence, or limited morphological differentiation, complicating both taxonomic resolution and subsequent research that depends on accurate species identification (Struck *et al*. 2018).

Historically, host specificity served as the primary criterion to distinguish groups of phytopathogenic strains within FOSC. Although FOSC displays a broad host range, individual pathogenic isolates typically infect only one or a few plant species (van Dam *et al*. 2016, Edel-Hermann & Lecomte 2019). Strains infecting the same host are grouped into a *forma specialis* (f. sp.), a nomenclatural system that links a strain’s name to its host specificity. For example, strains grouped into *F. oxysporum* f. sp. *lycopersici* can infect tomatoes, while those infecting chickpea are grouped into f. sp. *ciceris*. However, the *forma specialis* system has proven inadequate for defining taxonomic boundaries within the FOSC. First, it excludes the large number of putatively non-phytopathogenic strains that are part of FOSC and relies on incomplete host-range testing, which can lead to inconsistent or overlapping designations (Edel-Hermann & Lecomte 2019). Consequently, some strains are assigned to multiple *formae speciales*, such as f. sp. *apii* isolates capable of infecting several hosts including eggplant and asparagus, thereby overlapping with those belonging to *formae speciales melongenae* and *asparagi,* respectively (Epstein *et al*. 2017, Edel-Hermann & Lecomte 2019). Second and most critically, host specificity in FOSC has now been recognized as a polyphyletic trait, invalidating its use as a stable taxonomic marker (van Dam *et al*. 2016, 2017).

Given the limitations of the *forma specialis* system, several alternative frameworks have been explored for defining species boundaries in the FOSC. The biological species concept, based on reproductive isolation, has limited applicability because a sexual morph has not (yet) been observed in *F. oxysporum*, neither in nature nor under laboratory conditions (Lombard *et al*. 2019). Nonetheless, recent genomic studies have started to indicate that recombination might occur within certain populations (Laurence *et al*. 2014, Brankovics *et al*. 2017, McTaggart *et al*. 2021, Fayyaz *et al*. 2023). However, estimates of the number of recombining or independently evolving lineages vary widely, ranging from as few as two to as many as 21, depending on the analyzed genes and the applied delimitation method (Laurence *et al*. 2014, Brankovics *et al*. 2017, Lombard *et al*. 2019, Achari *et al*. 2020, McTaggart *et al*. 2021). Hence, species delineation based on reproductive isolates has not been successfully and comprehensively implemented in the FOSC to date.

The phylogenetic species concept offers a complementary perspective by defining species as the smallest monophyletic clusters of individuals that can be consistently recovered (Cracraft 1983, McKitrick & Zink 1988). Early multigene phylogenies divided the FOSC into three primary clades, a structure repeatedly supported using diverse genetic markers, including partial translation elongation factor 1-alpha (*tef1* / *EF-1α)*, entire nuclear ribosomal intergenic spacer (IGS), and mitochondrial small subunit rDNA (mtSSU) sequences (O’Donnell *et al*. 1998, 2004, Baayen *et al*. 2000). Subsequent studies using additional loci such as partial calmodulin (*cmdA* / *CAM*), RNA polymerase II largest subunit (*rpb1*), RNA polymerase II second largest subunit (*rpb2*), topoisomerase I (*Top1*), translation elongation factor 3 (*tef3*) and beta-tubulin (*tub2*) sequences have proposed five to eight clades, depending on analytical resolution and sampling scope (Laurence *et al*. 2011, Lombard *et al*. 2019, Achari *et al*. 2020, Nephalela-Mavhunga *et al*. 2021). Recent years have seen the description of numerous additional taxa, 11 new species since Lombard *et al*. (2019), bringing the current total to 51 described species within the complex (source: https://www.fusarium.org/). However, new species are frequently described based on under-sampled datasets that do not reflect the previously described diversity within the FOSC or based on very little sequence divergence (Zhang *et al*. 2023, Tan & Conroy 2024). This continuous expansion reflects both genuine evolutionary diversity in FOSC and the methodological challenges associated with the recognition of species. Furthermore, cultures of newly described species are often not available due to regulations under the Nagoya Protocol, or not all of the required genetic markers were sequenced, complicating comparisons of existing and newly described species. Collectively, the boundaries between species, populations, and pathotypes in FOSC therefore remain blurred, rendering FOSC taxonomy confusing and difficult to use. To address these challenges, we here used maximum likelihood- and coalescence-based analyses of nearly 2500 conserved marker genes obtained from around 660 FOSC genome assemblies, including newly generated whole-genome assemblies of 22 ex-type strains, representing almost half of the currently accepted species within FOSC, to generate a robust phylogeny of FOSC. Our revised phylogenomic framework provides a stable foundation for FOSC taxonomy and offers a critical resource for the broader research community.

## MATERIALS AND METHODS

### Constructing a FOSC phylogeny using commonly used marker genes

To construct a phylogeny of the FOSC using six commonly used marker gene sequences [*tef1*, *rpb1*, *rpb2*, *CAM*, *tub2*, and the internal transcribed spacer regions of the nrDNA (ITS)], we downloaded all available data from fusarium.org (Crous *et al*. 2021b) on 21-10-2025, yielding a total of 290 isolates, including 49 type isolates, as well as *Fusicolla matuoi* CBS 581.78 as an outgroup. For each marker gene, multiple sequence alignments were constructed with MAFFT v. 7.505 (Katoh & Standley 2013), using default settings, and trimmed with trimAl (Capella-Gutiérrez *et al*. 2009), using default settings. The single-gene alignments were then concatenated using the *create_concat* function from PhyKit (Steenwyk *et al*. 2021), and the resulting superalignment was used to construct a combined marker gene phylogeny of the FOSC, using IQ-TREE v. 3.0.1 using model finder plus for model selection (-m MFP) and 10000 UltraFast bootstraps (-B 10000) (Kalyaanamoorthy *et al*. 2017, Hoang *et al*. 2018, Wong *et al*. 2026). All trees were visualized using iTOL (Letunic & Bork 2024).

To calculate the number of parsimony informative sites for FOSC members, we removed alignment positions consisting entirely of gaps once we removed the outgroup using trimAl (-gt 0.01), and all N’s were subsequently replaced with gaps (Capella-Gutiérrez *et al*. 2009). The resulting alignments were used as input for the *parsimony_informative_sites* function from the PhyKit package (Steenwyk *et al*. 2021).

### Generating whole-genome assemblies of ex-types covering the diversity of the FOSC

To gain a higher resolution insight into the phylogenetic relationships within the FOSC, we generated *de novo* whole-genome assemblies for 48 isolates from the Westerdijk Fungal Biodiversity Institute’s biobank, including 22 ex-type isolates that can be used to place currently described species (Sup. Table 1). These isolates were selected to cover the full diversity of the marker-gene tree while including as many ex-type strains as possible for which we had access to.

Genomic DNA of all isolates was obtained from fungal mycelium grown on malt extract agar (MEA; Crous et al. 2019) plates using a hexadecyltrimethyl ammonium bromide (CTAB)-derived protocol adapted from Damm *et al*. (2008). The mycelium was scraped from the plates and added to 2 mL Eppendorf tubes which were stored at -80 °C before further processing. Before freeze-drying the mycelium, the Eppendorf tubes were opened, after which the opening of the tube was covered with Parafilm and some small holes were made in the Parafilm. The tubes were subsequently added to 50 mL tubes covered with Miracloth (Millipore, 475855-1R) and the mycelium freeze-dried in a Christ alpha1-4 LSC basic freeze dryer according to the manufacturer’s instructions. Three sterile metal ball bearings (3 mm diam.) were added to each 2 mL Eppendorf tube. The tubes were stored in -80 °C till further use. Starting the extraction protocol, the tubes were placed in liquid nitrogen. The frozen mycelium was then ground in a Qiagen TissueLyser III using a frequency of 30 Hz for 1 min. Preheated (to 65 °C) CTAB solution was added to each Eppendorf tube and the contents mixed by inversion, after which it was incubated in a heating block (Labnet Accublock) at 65 °C for at least 15 min, with manual inversion or shaking every 5 min. Three μL RNAse cocktail (Invitrogen AM2286) was added and the mix was incubated for 15 min at 37 °C. Working in a fumehood, 600 µL of chloroform : isoamyl alcohol (24:1) was added and the Eppendorf tubes shaken by hand for 5 min until the solution became milky. The fungal matrix was spun down for 10 min at 14000 rpm at 4 °C after which the aqueous phase was transferred into a new 2 mL centrifuge tube. Ice-cold 600 μL isopropanol was added to the supernatant and gently inverted to mix the contents. This mixture was spun down for 5 min at 14000 rpm at 4 °C after which the supernatant was discarded. The resulting pellet was washed with 70 % ethanol by adding 600 µL of cold 70 % ethanol to the centrifuge tube, repeating the 5 min centrifugation at 14000 rpm at 4 °C and discarding the supernatant. Any remaining visible ethanol was removed using a pipet. This washing step was repeated once. The DNA pellet was air-dried overnight or at 50 °C in a heating block for a maximum of 15 min and then resuspended in 50 μL DNAse-free ddH2O overnight at 10 °C. Nanodrop one (Thermo Fisher) and Qbit4 fluorometer (Invitrogen) were used to measure the DNA concentration and 260/280 ratio, and an agarose gel (1 % agarose, TAE buffer, 100 V, 45 min) was run to check for shearing of the genomic DNA.

Oxford Nanopore Technologies (ONT) raw reads were generated for 14 out of 48 isolates using a FLO-PRO114M Flowcell on a PromethION24 sequencer (ONT, UK) with SQK-NBD114-24 kit and acquired with MinKNOW v. 23.11.7. High-accuracy basecalling was enabled on the PromethION device with a minimum read length cutoff of 200 bases and a minimum Q score of 9. Reads passing quality filters were processed with Porechop v. 0.2.4 (https://github.com/rrwick/Porechop) to remove adapter sequences. Subsequently, reads longer than 1,000 bp and with a minimum quality score greater than Q10 were retained using NanoFilt v 2.6.0 (De Coster & Rademakers 2023) for downstream assembly. Summary statistics for raw, cleaned, and filtered datasets were generated using SeqKit v. 0.10.2 (Shen *et al*. 2016) . Genome size estimation was performed using a k-mer–based approach with Jellyfish v. 2.3.0 (Marçais & Kingsford 2011) and GenomeScope v. 2.0.0 (Ranallo-Benavidez *et al*. 2020), based on the filtered reads (Q10, length ≥1 kb). *De novo* genome assembly was carried out using Flye v. 2.9 (Kolmogorov *et al*. 2019) with scaffolding enabled and three iterations of polishing, leaving all other parameters set at default. Contigs shorter than 1,000 bp were removed from the final assembly. Assembly quality metrics were assessed using QUAST v. 5.0.2 (Gurevich *et al*. 2013), and genome completeness was evaluated with BUSCO v. 5.4.7 (Seppey *et al*. 2019, Tegenfeldt *et al*. 2025) using the Fungi and Eukaryota lineage datasets.

The remaining 34 isolates were sequenced using paired-end Illumina by BGI. The obtained reads were quality-filtered and trimmed using Trimmomatic v. 0.39 (Bolger *et al*. 2014) with: removal of leading and trailing low-quality or N bases (*LEADING:3* and *TRAILING:3*), 4-base sliding-window trimming when average quality dropped below a Phred score of 15 (*SLIDINGWINDOW:4:15*), removal of Illumina adapter sequences (*ILLUMINACLIP*) and discarding reads shorter than 70bp (*MINLEN:70*). The cleaned reads were assembled de novo using Velvet v. 1.2.10 (Zerbino & Birney 2008) with a k-mer size of 115 (-kmer 115). Assembly contiguity was evaluated using the *stats.sh* script from BBTools (Bushnell 2014), assessing total assembly size, scaffold count and N50. Additionally, BUSCO analysis was performed following the same protocol as for the ONT-based assemblies.

### Composing a whole-genome assembly dataset with public data

To place our newly sequenced *F. oxysporum* isolates into their genetic context, we assembled a diverse set of *F. oxysporum* isolates by downloading all 777 publicly available *F. oxysporum* whole-genome assemblies from NCBI GenBank on 15 July 2025. Genome completeness was assessed using Quast (v. 5.0.2) (Gurevich *et al*. 2013) and BUSCO (v. 6.0.0) (Tegenfeldt *et al*. 2025) with default settings, the latter using the *Nectriaceae* dataset for the BUSCO marker set. Duplicate genome assemblies originating from the same isolate were then manually removed, retaining the genome assembly with the highest single-copy BUSCO completeness or, in cases of equal completeness between the most complete assemblies, the one with the fewest contigs. Assemblies with a single-copy BUSCO completeness below 97 % were excluded to ensure high assembly quality. Additionally, we removed assemblies GCA_016802495.1, GCA_016802205.1, and GCA_016802225.1, as these originated from isolates that were *in silico* Minimum Information about a Single Amplified Genome (MISAG) hybrids, generated by imputing sequencing data onto the reference genome of *Fo*4287. Assemblies GCA_01959348.1 and GCA_0147702115.1 were likewise removed, as they were marked as originating from unverified source organisms and clustered with the outgroup during preliminary phylogenetic analysis. This resulted in a final set of 612 whole-genome assemblies from distinct *F. oxysporum* isolates used in this work (Sup. Table 2). These publicly available assemblies were then combined with our newly sequenced isolates, yielding a final dataset of 660 whole-genome assemblies of FOSC isolates, including 22 ex-type isolates. The genome assemblies of three *Fusarium redolens* isolates (GCA_017562085.1, GCA_020744475.1, and GCA_035773055.1) were included as an outgroup for subsequent analyses where appropriate (Sup. Table 2).

### Constructing a FOSC phylogeny using the *Fusarium oxysporum* core genome

#### Obtaining single-gene multiple sequence alignments of BUSCO genes

To extract all BUSCO genes shared in single copy across all isolates in our dataset, we used ArboPhyl v. 1 (https://github.com/WesterdijkInstitute/ArboPhyl). Amino-acid multiple sequence alignments (MSAs) were constructed for each conserved BUSCO gene using MAFFT v. 7.505 in high-speed mode (Katoh & Standley 2013). Corresponding nucleotide sequences for each gene were threaded back onto the amino acid alignments using PAL2NAL v. 14 with the *nogap* setting, trimming all gaps from the alignments (Suyama *et al*. 2006). IQ-TREE v. 2.1.4_beta was used to compute a bootstrapped gene tree per nucleotide MSA, using ModelFinderPlus (-m MFP) and 10,000 UltraFast bootstraps (-B 10000) (Kalyaanamoorthy *et al*. 2017, Hoang *et al*. 2018, Minh *et al*. 2020).

#### Obtaining maximum-likelihood- and coalescence-based species trees

To obtain a coalescence-based species phylogeny, we used wASTRAL v. 1.23.3.7, part of the ASTER package, with the gene trees in hybrid weighting mode (--mode 1), 16 rounds of subsampling per exploration step (-s 16), 16 rounds of initial placement (-r 16), and outputting detailed support information (-u 2) (Zhang & Mirarab 2022, Zhang *et al*. 2025). Since wATRAL returns a tree with branch lengths in the non-standard unit of coalescent units and support values as local posterior probabilities, we used a custom script to remove all branch length and node support information from the species tree, leaving only an unrooted topology. IQ-TREE v. 3.0.1 was then used with the ASTRAL topology as a constraint tree (-g) to calculate branch lengths in Substitution Units (SU) and UltraFast bootstrap support (-B 10000), using the *keep-ident* setting and supplying the previously computed evolution model per gene using a partition file (-p) (Chernomor *et al*. 2016, Hoang *et al*. 2018, Wong *et al*. 2026). Gene concordance factors (gCF) were subsequently added to this tree using the *gcf* setting, with the individual gene trees as inputs (Mo *et al*. 2023). Finally, site concordance factors (sCF) were added to the tree using the *scfl* setting with the trimmed nucleotide gene MSAs as input (Mo *et al*. 2023). This yielded a species tree where the topology of the tree was computed using wASTRAL and the branch lengths, bootstrap-, gCF-, and sCF-support values were computed with IQ-TREE. This ATRAL-based tree is hereafter referred to as the “*coalescence*-based species tree”.

To cross-validate the coalescence tree topology and reveal any strong methodological biases, we also computed a maximum likelihood-based species tree with IQ-TREE v. 3.0.1, with a partition file (-p) specifying previously computed substitution models for each gene, and 10000 UltraFast bootstrap computations (-B 10000) (Hoang *et al*. 2018, Wong *et al*. 2026). As for the coalescence-based species tree, gCF, and sCF support values were subsequently added to this maximum-likelihood-based species tree using the same parameters described above. This tree is hereafter called the “*maximum likelihood*-based species tree”.

We then identified robust clusters of isolates forming monophyletic clades in both the coalescense-and maximum likelihood-based species trees. These groups were then annotated as *putative species*” and used throughout subsequent analyses.

#### Phylogenetic analyses of subset data

To further assess the robustness of our phylogenetic analyses and to assess whether tree topology is driven by a particular subset of marker genes, we subsampled our dataset and recomputed phylogenetic trees per subsample. We first computed, for each BUSCO gene, the percentage of parsimony informative sites, defined as a site that has at least two different nucleotides that occur at least twice across all included isolates, using the *parsimony_informative_sites* function from Phykit v. 2.0.3 (Steenwyk *et al*. 2021). We then generated 19 different subsets of gene trees: (i) a subset excluding the top and bottom 10 % of genes in terms of percentage of parsimony informative sites (“80-percent subset”); (ii) nine subsets of 277 randomly selected genes (total number of BUSCO genes divided by nine), with each gene included in exactly one subset (“random subsets”); and (iii) nine subsets of 277 genes ordered by their percentage of parsimony-informative sites, with each gene included in exactly one subset (“ordered subsets”). For each subset, we inferred phylogenetic trees using both wASTRAL and IQ-TREE, as described above. These subset phylogenetic trees were then visualized using iTOL (Letunic & Bork 2024) and manually assessed for their recovery of the delineated putative species as monophyletic lineages, as well as their relative positions as compared to the *maximum likelihood*-based species tree.

### Assessing recombination and introgression with SplitsTree

A possible explanation for the variation observed among the species trees is exchange of genetic material through hybridization, recombination or introgression. To investigate this possibility, we used SplitsTree v. 6.5.1 to compute a SplitsNetwork based on a concatenation of all 2493 nucleotide MSAs (Huson & Bryant 2024). Clusters were then manually annotated according to the putative species identified. For each putative species, trimmed nucleotide MSAs were subsampled by removing all sequences from strains that were not included in that putative species. These cluster-specific MSAs were then concatenated and used for cluster-specific SplitsTree analyses.

### Recovering phylogenetic species with gene-independent variation

To further corroborate the putative species, we defined core genomes for all FOSC isolates, excluding accessory genomic regions that are known to be horizontally transferred between FOSC isolates (Ma *et al*. 2010) and could otherwise introduce conflicting phylogenomic signal. Core genomic regions were extracted from all 657 FOSC genome assemblies; this dataset excluded the three *F. redolens* outgroup isolates (GCA_020744475.1, GCA_035773055.1, and GCA_017562085.1) and three isolates that consistently clustered outside of the FOSC in our analyses (GCA_014324465.1, GCA_019157295.1, and GCA_024865645.1).

We first reconstructed the core genome of *F. oxysporum* f. sp. *lycopersici* (*Fo*4287), a genome assembly that has been extensively studied in the past (Ma *et al*. 2010). We downloaded the *Fo*4287 assembly (GCA_000149955.2), and manually removed accessory chromosomes 3, 6, 14 and 15, as well as all supercontigs included in the assembly. To remove accessory regions at the chromosome ends of chromosomes 1 and 2, we aligned the corresponding supercontigs 27 (DS231722) and 31 (DS231726), respectively, and then removed the aligned accessory regions from those core chromosomes. Subsequently, we aligned each of the 657 FOSC genomes this *Fo*4287 ‘core-genome’ using minimap2 v. 2.30-r1287 with the preset for ∼0.1 % sequence divergence (-x asm5). We then used a custom script to select for each FOSC isolate contigs of at least 10 kb with combined non-overlapping query coverage of at least 70 %, and generated core-genome assemblies from these.

Using these core-genome assemblies, we next calculated pairwise nucleotide identity by running reciprocal minimap2 v. 2.30-r1287 searches, using the *-x asm5* setting as well as the long CS tag (--cs=long). The resulting mappings were sorted in genomic order relative to the reference genome and subsequently used for the paftools.js call function (part of the minimap2 suite), which calls variants between the reference and the query genome, using default settings. We subsequently calculated nucleotide identity as (1 – number of substitutions / bases covered) * 100. As reciprocal searches did not necessarily yield identical nucleotide identity values, owing to slight differences between minimap2 alignments, we calculated final nucleotide identity between two genomes as the average of their reciprocal comparisons. Finally, we clustered the dataset based on their pairwise nucleotide similarity using UPGMA; distances were calculated as 100 - percent identity. The resulting dendrogram and heatmap were visualized using the seaborn package in Python.

To further assess genetic diversity, we constructed a single nucleotide polymorphism (SNP) matrix of all our FOSC core-genome assemblies. The SNPs were called relative to the *Fo*4287 core-genome described above, using minimap2 and paftools. The resulting SNP data were integrated using bcftools merge and then filtered for biallelic SNPs using bcftools view (-v snps - m2 -M2). In the resulting matrix, all positions not called as variants were treated as equal to the reference. Each SNP in the matrix was re-coded such that the dominant allele was represented as “0” instead of “1”, following best practices for SNP-based PCA analyses (Gauch Jr *et al*. 2019). To remove uninformative SNPs, the matrix was filtered for a minimal allele frequency of 0.5 %. We then calculated a distance matrix using Hamming distance and performed PCoA as implemented in Python scikit-bio. To gain further insight into the difference between isolates within clades 2 and 3, this workflow was repeated after removing isolates in Clade 1.

### Investigating current marker efficacy for recapturing putative species

To assess the ability of the six commonly used sequenced marker genes for FOSC isolates (Table 1) to recapture the putative species delineated in our work, we extracted these marker gene sequences from all the 663 whole genome assemblies in our dataset. We used BLASTn v. 2.12.0+ (Camacho *et al*. 2009) with a single maximum target sequence (-max_target_seqs 1) and a single maximum High-scoring Segment Pair (-max_hsps 1) to extract sequences for *tef1, rpb2, rpb1, CAM, tub2,* and *ITS* for all 663 assemblies in the whole-genome dataset, using FOSC sequences CBS 148206 for *tef1*, CBS 144744 for *rpb2*, GUCC_190024.2 for *rpb1*, CBS 148222 for *ITS*, CBS 413.90 for *tub2* and LLC3505 for *CAM* as queries (Sup. Table 3). These query sequences were selected from the fusarium.org database based on maximum gene length and minimum number of N’s. As described previously for the original marker-gene tree, single-gene multiple sequence alignments were constructed using MAFFT v. 7.505 on standard settings (Katoh & Standley 2013), and used as input for IQ-TREE v. 3.0.1 running a partition file (-p), Model Finder Plus per partition (-m MFP) and 10000 UltraFast bootstrap replicates (-B 10000) (Kalyaanamoorthy *et al*. 2017, Hoang *et al*. 2018, Wong *et al*. 2026). The resulting tree was visualized in iTOL (Letunic & Bork 2024), and the putative species definitions determined above were mapped onto this tree.

### Finding alternative marker gene sequences for species identification

To identify BUSCO genes that could be used as additional marker genes to delineate species within FOSC, we supplemented phylogenies built solely from *tef1* and *rpb2,* as these two marker genes are the most commonly sequenced, and contain the highest number of informative residues (Table 1), with additional markers used here. We used a custom script to screen each of the 2493 BUSCO genes for unique positions conserved within a putative species, but distinct from all other isolates. Starting from *tef1* and *rpb2*, we then identified the minimal combination of genes containing at least one unique position per putative species. To confirm that this selected set of traditional and BUSCO marker genes could recapitulate all putative species described here as monophyletic clades, we constructed a phylogenetic tree using the same protocol as described above for the construction of the maximum likelihood species tree.

## RESULTS AND DISCUSSION

### Current marker genes contain insufficient information for species delineation in the Fusarium oxysporum species complex

Phylogenetic analyses of members of the FOSC commonly rely on variable combinations of marker genes such as *tef1*, *rpb2*, *rpb1*, *CAM*, ITS, and *tub2* (O’Donnell *et al*. 1998, 2004, Baayen *et al*. 2000, Laurence *et al*. 2011, 2014, Brankovics *et al*. 2017, Lombard *et al*. 2019). To assess the capacity of these marker genes to robustly delineate species within FOSC, we first collected nucleotide sequences for a total of 287 putative FOSC isolates, together with outgroup isolate *Fusicolla matuoi* CBS 581.78. After exploratory analysis, the ex-types for *Fusarium kamalianum* (NFCCI 5154) and *Fusarium nashariae* (BRIP 65609b) were excluded, as they consistently clustered outside the FOSC and closer to the outgroup. Notably, sequence availability varied considerably among markers: *tef1* and *rpb2* were most widely represented (286 and 272 sequences, respectively), whereas ITS and *tub2* were available for fewer isolates (109 and 110 sequences, respectively) (Table 1). Although the ingroup-only alignment of *rpb1* was approximately twice as long as the others, it contained relatively few (5.3 %) parsimony informative sites, while 16 % of all sites of *tef1* are parsimony informative.

**Table 1.**
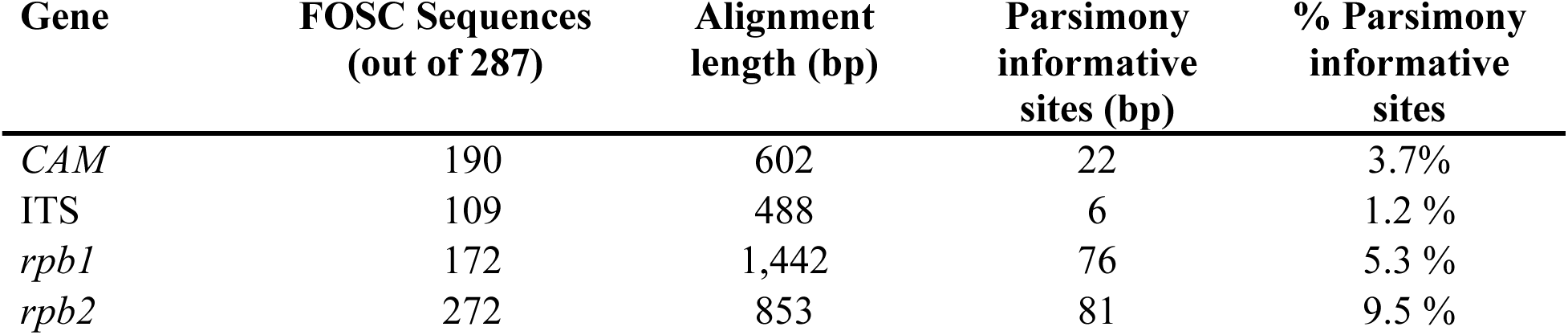
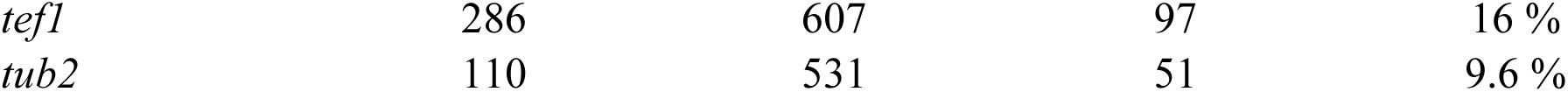
Marker genes differ in their abundance, length, and proportion of parsimony informative sites. Metadata of marker genes extracted from fusarium.org. For the calculation of alignment lengths, parsimony informative sites, and % parsimony informative sites, we first removed the outgroup isolate *Fusicolla matuoi* CBS 581.78 and subsequently removed all gap-positions created by the removal of this sequence.

We then inferred a maximum-likelihood phylogenetic tree of 287 FOSC isolates based on the concatenated alignment of *tef1*, *rpb2*, *rpb1*, *CAM*, ITS, and *tub2*; bootstrap support values ranged from 2–100 %, with most basal nodes having below 95 % ultrafast bootstrap support and thus being poorly supported. Notably, this phylogenetic tree did not recover the three clades commonly described within the FOSC (O’Donnell *et al*. 1998, 2004, Baayen *et al*. 2000, Brankovics *et al*. 2017, van Dam *et al*. 2017, McTaggart *et al*. 2021) (Fig. 1, Sup. Fig. 1). The canonical Clade 3 did not form a monophyletic clade but instead was split into three separate clades, two of which appear closely related to the outgroup. Moreover, Clade 1, normally recovered as the first diverging clade within the FOSC, occupied a position between clades 2 and 3. Furthermore, several non-type isolates clustered closer to type isolates of other species rather than with the type isolate of their own designated species, such as observed for *F. odoratissimum*, *F. veterinarium*, or *F. fabacearum*, where multiple ex-types and species names were intermixed within larger clades. This variable placement likely reflects two underlying issues of misidentification: first, some isolates may have originally been identified using trees that lacked a comprehensive set of ex-types, meaning the correct species label could never have been assigned in the first place; and second, correct identification requires the appropriate combination of marker genes, so isolates classified using insufficient or conflicting phylogenetic signal, may have been mislabeled. Finally, several ex-type strains were separated by only few substitutions or, in some cases, were more closely related to each other than to any other isolate, as exemplified by the type isolates *F. callistephi* CBS 187.53 and *F. inflexum* CBS 716.74. Taken together, this suggests that these six markers are insufficient to robustly delineate and support the currently accepted species within FOSC (Sup. Fig. 2).

**Fig. 1.**
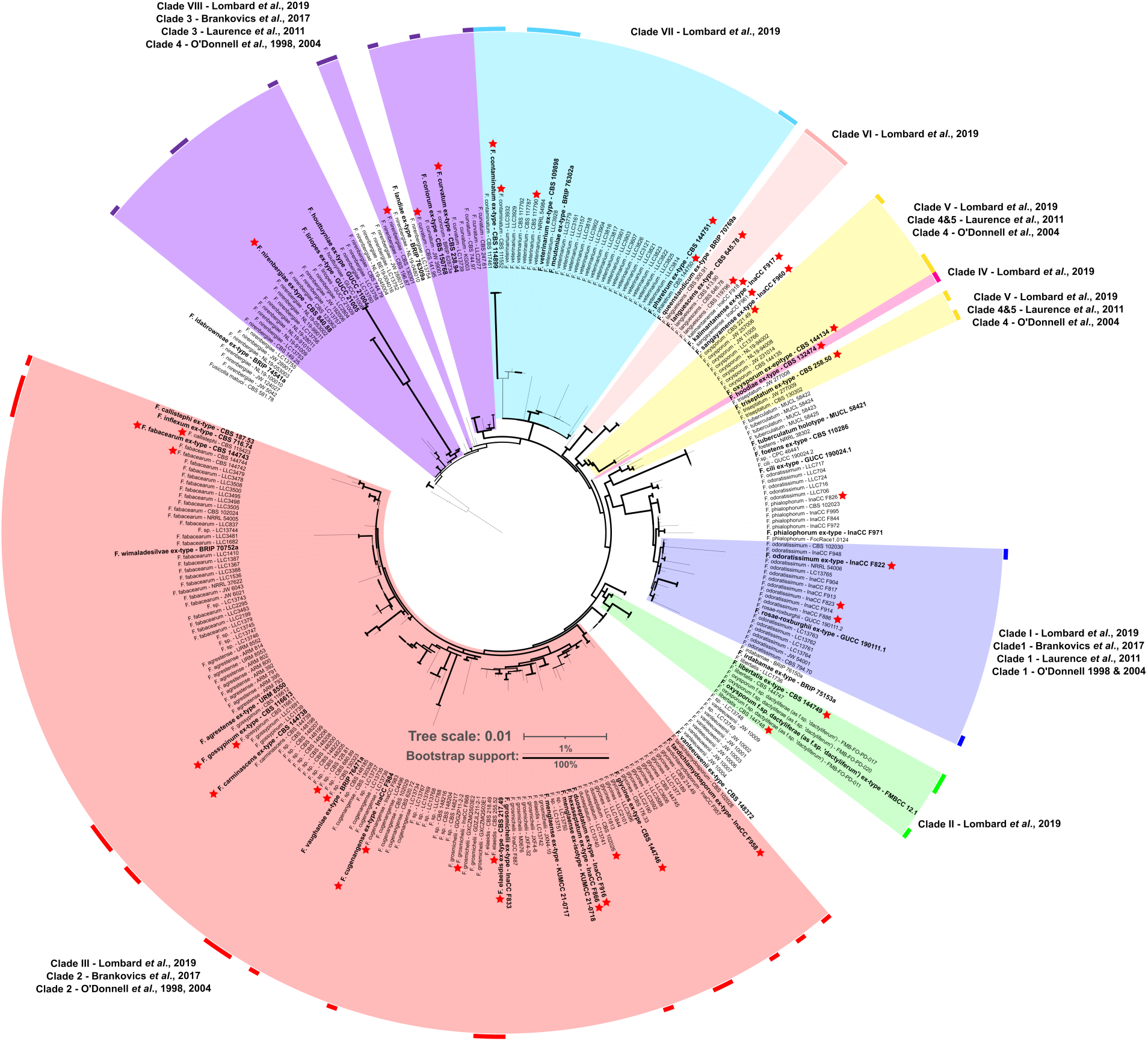
A phylogenetic tree based on six commonly used marker sequences is insufficient to robustly delineate species in the *Fusarium oxysporum* species complex. The maximum-likelihood based phylogenetic tree was inferred using a concatenated nucleotide alignment of *tef1, rpb2, rpb1, CAM*, ITS, and *tub2* obtained from 287 FOSC isolates and outgroup isolate *Fusicolla matuoi* CBS 581.78 from fusarium.org. Type isolates are highlighted in bold, and isolates selected for whole-genome sequencing (Fig. 2) are emphasized with a red star. Branch length represents substitution per sites; branch thickness indicates ultrafast bootstrap support, and the outgroup branch was shortened for visualization purposes. The outer ribbon indicates isolates with a clade annotation from previous studies. Background shadings indicate monophyletic species extrapolated from isolates with a clade annotation. Labels next to background shadings indicate names given to clades in previous studies.

### Whole-genome sequencing and assembly of diverse FOSC isolates to obtain additional markers for species delineation

To leverage an extended set of gene markers for robust species delineation within FOSC, we selected 48 isolates that span the phylogenetic diversity of the species complex for whole-genome sequencing (Fig. 1). This selection included 24 ex-type isolates, representing, to our knowledge, the largest set of FOSC ex-types sequenced to whole-genome level to date. Isolates were sequenced using Nanopore sequencing and assembled using Flye (Kolmogorov *et al*. 2019). We observed considerable differences in the size and the number of contigs, ranging from 43 to 57 Mb and between 12 to 4080 contigs (Sup. Table 1, Fig. 2), with N50 ranging between 0.2 and 5.0 Mb. Given that FOSC isolates typically have 11 core chromosomes (Ma *et al*. 2010), we expect a subset of these assemblies to approach chromosome-level contiguity. Despite the variation in genome size and fragmentation, all assemblies captured most of the gene space, as reflected by consistently high single-copy BUSCO completeness scores against the Nectriaceae dataset, ranging from 98.9 to 99.5 % across all assemblies (Sup. Table 1, Fig. 2). These single-copy BUSCO marker genes thus represent an excellent starting point for phylogenetic analyses of the FOSC.

**Fig. 2.**
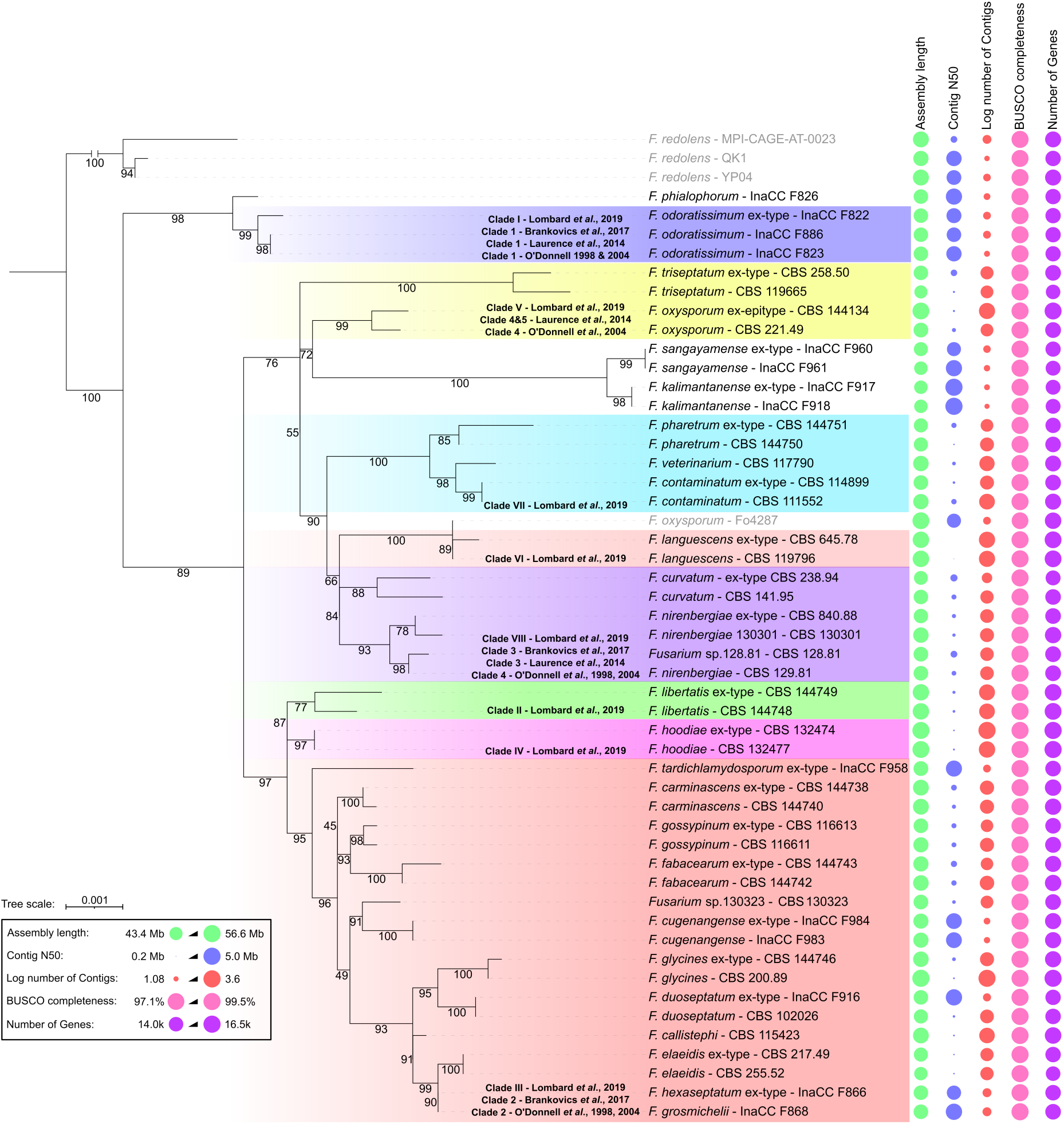
Newly generated whole-genome assemblies of FOSC isolates, including 22 ex-types, are of high quality and completeness. Whole-genome sequencing and assembly of 48 *Fusarium oxysporum* isolates, including 22 ex-types. Assembly length (genome size in bp), contig N50 and log number of contigs is shown. Completeness of the gene space is indicated using the total number of predicted protein-coding genes as well as BUSCO completeness; *F. oxysporum* isolate *F. oxysporum* f.sp. *lycopersicum* 4287 is shown as reference (Ma *et al*. 2010). The maximum-likelihood phylogenetic tree is based on the concatenated alignments of the *tef1, rpb2, rpb1, CAM*, ITS, and *tub2* marker genes extracted directly from the genome assemblies to emphasize the diversity of *F. oxysporum* ex-types selected for whole–genome sequencing (see also Fig. 1). Background shading indicates the previously described clades, following Fig. 1.

### A deeply sampled phylogenetic tree of the FOSC using whole-genome data

Previous attempts to resolve the phylogenetic relationship of ex-types in FOSC and to robustly delineate species within this complex mostly relied on a limited set of marker genes and/or restricted sampling of species and strains (O’Donnell *et al*. 1998, 2004, Baayen *et al*. 2000, Laurence *et al*. 2011, 2014, Brankovics *et al*. 2017, Lombard *et al*. 2019, Nephalela-Mavhunga *et al*. 2021). Here, we sought to overcome these limitations by combining the genome sequences of the ex-types generated in this study with publicly available genomic resources to construct a deeply sampled and robust phylogeny of FOSC. We first obtained 777 FOSC whole-genome assemblies from NCBI (accessed 15 July 2025). Single-copy BUSCO completeness across these assemblies ranged from 4.6 to 99.3 %, with 627 isolates reaching a single-copy completeness of at least 97 %. We excluded 15 genome assemblies: some came from strains already represented elsewhere in the dataset, some were MISAG assemblies imputed onto Fo4287, and others clustered outside the FOSC in preliminary analysis. This left 612 high-quality FOSC genome assemblies. Adding our 48 newly generated assemblies gave a final dataset of 660 FOSC isolates, including 22 ex-type isolates. We also added three *Fusarium redolens* assemblies as an outgroup (Geiser *et al*. 2021, Gomez-Chavarria *et al*. 2024), bringing the total to 663 genome assemblies (Sup. Table 2).

To obtain suitable markers for FOSC phylogenetics, we used single-copy BUSCO genes, identifying 2493 single copy BUSCO genes conserved across all 663 genome assemblies (100 % taxon occupancy). Visual inspection of the resulting multiple sequence alignments revealed that individual gene predictions occasionally included substantial additional up- or downstream sequences outside the coding regions, resulting in predicted sequences considerably longer than expected. To remove those regions prior to downstream analyses, we applied an aggressive trimming strategy that removed all alignment positions containing gaps in any of the 663 sequences. After trimming, we retained 2493 aligned BUSCO genes, with alignment lengths ranging from 168 to 6.456 nucleotides (mean: 1355.37; SD: 817.73), and the number of parsimony informative sites ranging between 5 and 781 (mean: 136.19; SD: 110.81) (Sup. Table 4). The percentage of parsimony informative sites lay between 1.5 and 88.1 % (mean: 10,07%; SD: 6.38). This indicates that a substantial number of these markers contain more informative sites than the gene markers commonly used previously (Table 1, Sup. Table 4). The complete concatenated gene-marker sequence is 3.378.942 nucleotides long, with a total of 339.521 parsimony informative sites (10.05%).

We then performed maximum-likelihood based phylogenetic analysis implemented in IQ-TREE, searching for optimal nucleotide substitution models for each gene partition. As expected, both the outgroup and the FOSC were recovered as monophyletic taxa, and most of the internodes received maximum ultra-fast bootstrap support (Fig. 3, Sup. Fig. 3); the ultra-fast bootstrap support values ranged between 44 % and 100 % overall, with lower values predominantly found near leaf nodes and among closely related isolates. Three isolates annotated in NCBI as FOSC consistently cluster outside of the FOSC in our analysis (GCA_014324465.1, GCA_019157295.1, and GCA_024865645.1). We therefore conclude that these isolates are most likely part of another *Fusarium* species complex or species, more closely related to the FOSC than *F. redolens*, such as *F. foetens, F. nisikadoi*, *F. fujikuroi* or *F. newnesense* (Schroers *et al*. 2004, Geiser *et al*. 2021, Gomez-Chavarria *et al*. 2024).

The three previously described clades (Clades 1, 2, and 3) were recognized in our analyses, where the phylogenomic distance separating Clade 1 from either of the two other clades was larger than those separating Clades 2 and 3. Additionally we recovered 14 well-supported phylogenetic lineages, each containing at least one ex-type, as well as two additional lineages lacking a previously described ex-type; we opted not to formally name these taxa, designating them “Undescribed 1” and “Undescribed 2” pending comparison against a more comprehensive set of ex-type isolates. Within these 16 delineated lineages, several lineages contained multiple ex-type isolates, for example the ex-type isolates of *F. kalimantanense* as well as *F. sangayamense* clustered within the same lineage (Fig. 3), implying that, under the assumption that these 16 lineages constitute genuine phylogenetic species, several previously described species would need to be synonymized (Table 2). Ten species can be synonymized with certainty, as their ex-type isolates were included in this analysis. Four additional isolates were included that were assigned to specific species in fusarium.org based on previous phylogenetic studies but were not indicated as having any (ex-)type status (the ex-type cultures of those species were not available to us for genome sequencing). For these species (*F. phialophorum*, *F. veterinarium*, *F. callistephi* and *F. grosmichelii*), we expect the ex-type isolates to cluster in similar positions in our phylogeny, and thus already propose to synonymize these names pending further analysis.

**Table 2.**
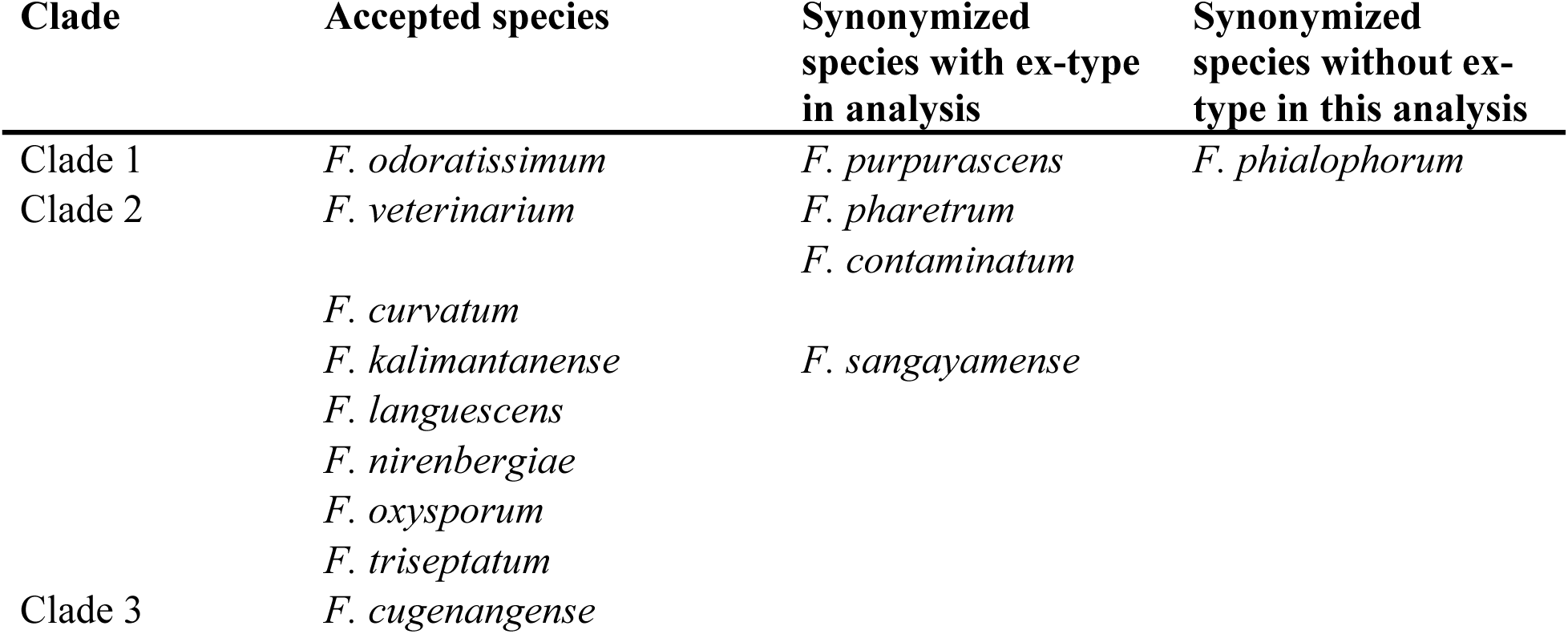
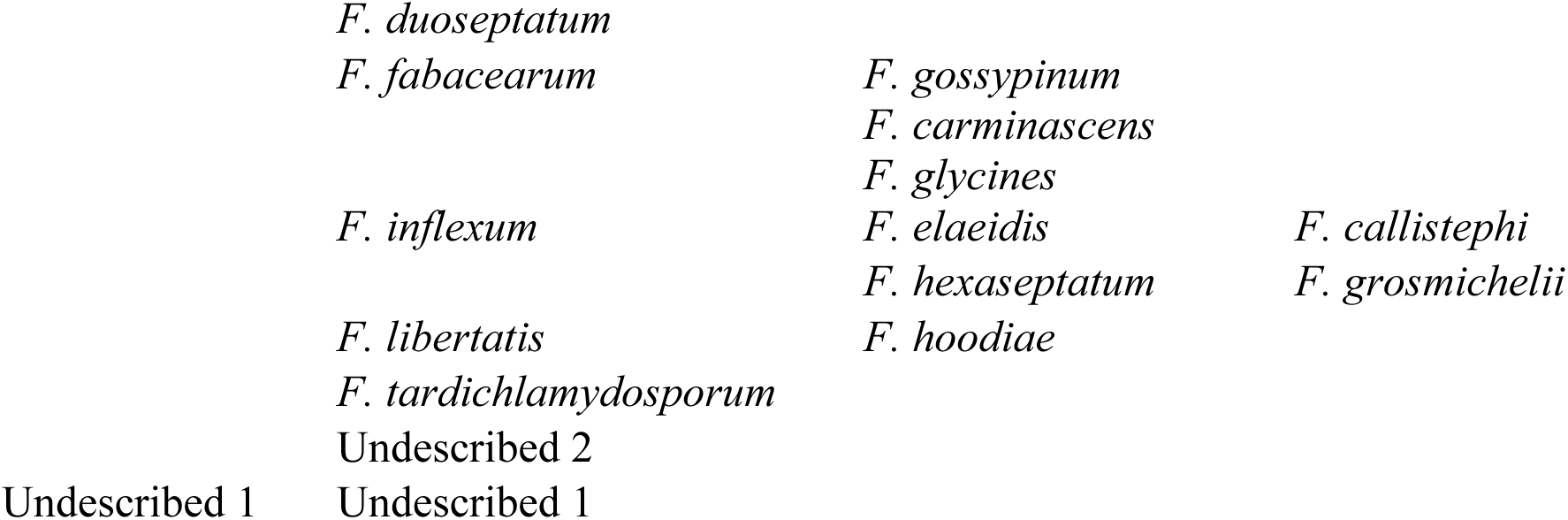
Overview of proposed synonymies of FOSC species included based on our phylogenomic analyses.

### Taxonomy

***Fusarium cugenangense*** Maryani *et al*., *Stud. Mycol*. **92**: 181. 2018. MB 826807.

***Fusarium curvatum*** L. Lombard & Crous, *Persoonia* **43**: 21. 2018. MB 826837.

***Fusarium duoseptatum*** Maryani *et al*., *Stud. Mycol*. **92**: 181. 2018. MB 826804.

***Fusarium fabacearum*** L. Lombard *et al*., *Persoonia* **43**: 24. 2018. MB 826839.

*Synonyms*: *Fusarium gossypinum* L. Lombard & Crous, *Persoonia* **43**: 26. 2018. MB 826841.

*Fusarium tardicrescens* Maryani *et al*., *Stud. Mycol*. **92**: 185. 2018. MB 826809. Nom. inval., Art. 40.7 (Shenzhen).

*Fusarium tardicrescens* Maryani *et al*., Persoonia **43**: 69. 2019. MB 828959.

*Fusarium carminascens* L. Lombard *et al*., *Persoonia* **43**: 19. 2018. MB 826835.

*Fusarium glycines* L. Lombard *et al*., *Persoonia* **43**: 25. 2018. MB 826840.

*Notes*: Of the names listed here, *F. fabacearum* is the most cited species in scientific literature.

*Fusarium tardicrescens* was reduced to synonymy under *F. fabacearum* by Lombard *et al*. (2022).

***Fusarium libertatis*** L. Lombard & Crous, *Persoonia* **43**: 29. 2018. MB 826844.

*Synonym*: *Fusarium hoodiae* L. Lombard *et al*., *Persoonia* **43**: 27. 2018. MB 826842.

*Note*: The name *F. libertatis* is the more frequently cited name in literature for this clade.

***Fusarium inflexum*** R. Schneid., *Phytopathol. Z*. **82**(1): 80. 1975. MB 314216.

*Synonyms*: *Fusarium grosmichelii* Maryani *et al*., *Stud. Mycol*. **92**: 176. 2018. MB 826803.

*Fusarium hexaseptatum* Maryani *et al*., *Stud. Mycol*. **92**: 183. 2018. MB 826808.

*Fusarium elaeidis* L. Lombard & Crous, *Persoonia* **43**: 23. 2018. MB 826838.

*Fusarium callistephi* L. Lombard & Crous, *Persoonia* **43**: 15. 2018. MB 826833.

*Note*: *Fusarium inflexum* was shown to be the older name for this clade by Han *et al*. (2025).

***Fusarium kalimantanense*** Maryani *et al*., Stud. Mycol. **92**: 187. 2018. MB 826810.

*Synonym*: *Fusarium sangayamense* Maryani *et al*., *Stud. Mycol*. **92**: 187. 2018. MB 826811.

***Fusarium languescens*** L. Lombard & Crous, *Persoonia* **43**: 28. 2018. MB 826843.

***Fusarium nirenbergiae*** L. Lombard & Crous, *Persoonia* 43: 29. 2018. MB 826845.

***Fusarium odoratissimum*** Maryani *et al*., *Stud. Mycol*. **92**: 159. 2018. MB 826800.

*Synonyms*: *Fusarium purpurascens* Maryani *et al*., *Stud. Mycol*. **92**: 160. 2018. MB 826801.

*Fusarium phialophorum* Maryani *et al*., *Stud. Mycol*. **92**: 169. 2018. MB 826802.

***Fusarium oxysporum*** Schltdl., *Flora Berolinensis, Pars secunda: Cryptogamia*: 139. 1824. MB 218372. See Fusarium.org for list of synonyms

***Fusarium tardichlamydosporum*** Maryani *et al*., *Stud. Mycol*. **92**: 181. 2018. MB 826805.

***Fusarium triseptatum*** L. Lombard & Crous, *Persoonia* **43**: 34. 2018. MB 826847.

***Fusarium veterinarium*** L. Lombard & Crous, *Persoonia* **43**: 35. 2018. MB 826849.

*Synonyms*: *Fusarium pharetrum* L. Lombard & Crous, *Persoonia* **43**: 32. 2018. MB 826846.

*Fusarium contaminatum* L. Lombard & Crous, *Persoonia* **43**: 20. 2018. MB 826836.

*Note*: The name *F. veterinarium* is the more frequently cited name in literature for this clade.

Our phylogeny and proposed species delineation is broadly concordant with prior studies at the clade level, with numerous putative species delineated here consistently recovered as monophyletic in earlier analyses that used independent datasets (Table 3) (Brankovics *et al*. 2017, van Dam *et al*. 2017, Lombard *et al*. 2019, Achari *et al*. 2020, McTaggart *et al*. 2021). For example, all putative species described here and represented in the ten-locus phylogeny of Brankovics *et al*. (2017) and the 441-gene phylogeny of van Dam *et al*. (2017) formed monophyletic clades in those analyses as well. The phylogeny of Lombard *et al*. (2019), who delineated eight clades and 21 species rather than the traditional three clades, was largely congruent with ours, aside from the proposed synonymies (Table 2). We only observed a few detailed differences. For example, Clade III isolates recovered by Lombard *et al*. (2019) are split among *F. cugenangense*, *F. duoseptatum*, *F. inflexum*, *F. fabacearum*, and *F. nirenbergiae* in our analysis, which we attribute to the limited phylogenetic signal in the original marker-gene dataset. Similarly, comparison with phylogenies proposed by Achari *et al*. (2020) and McTaggart *et al*. (2021), both of which are based on whole-genome assemblies, revealed only minor incongruences. Clade 4 described by Achari *et al*. (isolate RBG6505) falls within our Clade 2 as “Undescribed 2”, likely because their dataset lacked *F. tardichlamydosporum and F. libertatis*, which we recovered as early diverging lineages relative to “Undescribed 2”. Notably, the analyses by McTaggart *et al*. (2021) shared 327 isolates with our dataset and are thus most directly comparable. Of the 16 putative species delineated here, only *F. kalimantanense* was absent from their dataset. Their 12 phylogroups mapped directly onto our three clades (Phylogroup IX = Clade 1; Phylogroups I, IV, and XII = Clade 2; Phylogroups II, III, V, VI, VII, VIII, and XI = Clade 3), with isolate Fo65 (GenBank assembly GCA_014324465, genome assembly ASM1432446v1; their Phylogroup X) being the only exception. In both our analysis and McTaggart *et al*. (2021), this isolate was the first branching isolate of the ingroup. Since neither study included isolates from the *F. nisikadoi* or *F. newnesense* species complexes, which are more closely related to the FOSC than the outgroups used by both studies (Geiser *et al*. 2021, Gomez-Chavarria *et al*. 2024), we consider it likely that this isolate is misclassified as a FOSC isolate and is instead part of either *F. nisikadoi* or *F. newnesense* species complex. Both our study and McTaggart *et al*. (2021) also recovered isolate Fo24 (GenBank assembly GCA_014337855, genome assembly ASM1433785v1; “Undescribed 1” here; Phylogroup XI in McTaggart *et al*. 2021) in the same position, diverging prior to the split between Clades 2 and 3, consistent with McTaggart and colleagues’ divergence-time estimates (∼1.7 Mya for Clade 1, ∼0.93 Mya for Phylogroup XI, ∼0.90 Mya for the Clade 2/3 split), suggesting that this isolate may represent an additional lineage within the FOSC. Even though we could not directly compare our analyses with those of O’Donnell *et al*. (1998, 2004), Baayen *et al*. (2000), and Laurence *et al*. (2011, 2014), as these studies share little to no overlap with our dataset, the overall concurrence between our analyses and previous work suggests that the here defined taxonomic lineages are robust.

**Table 3.**
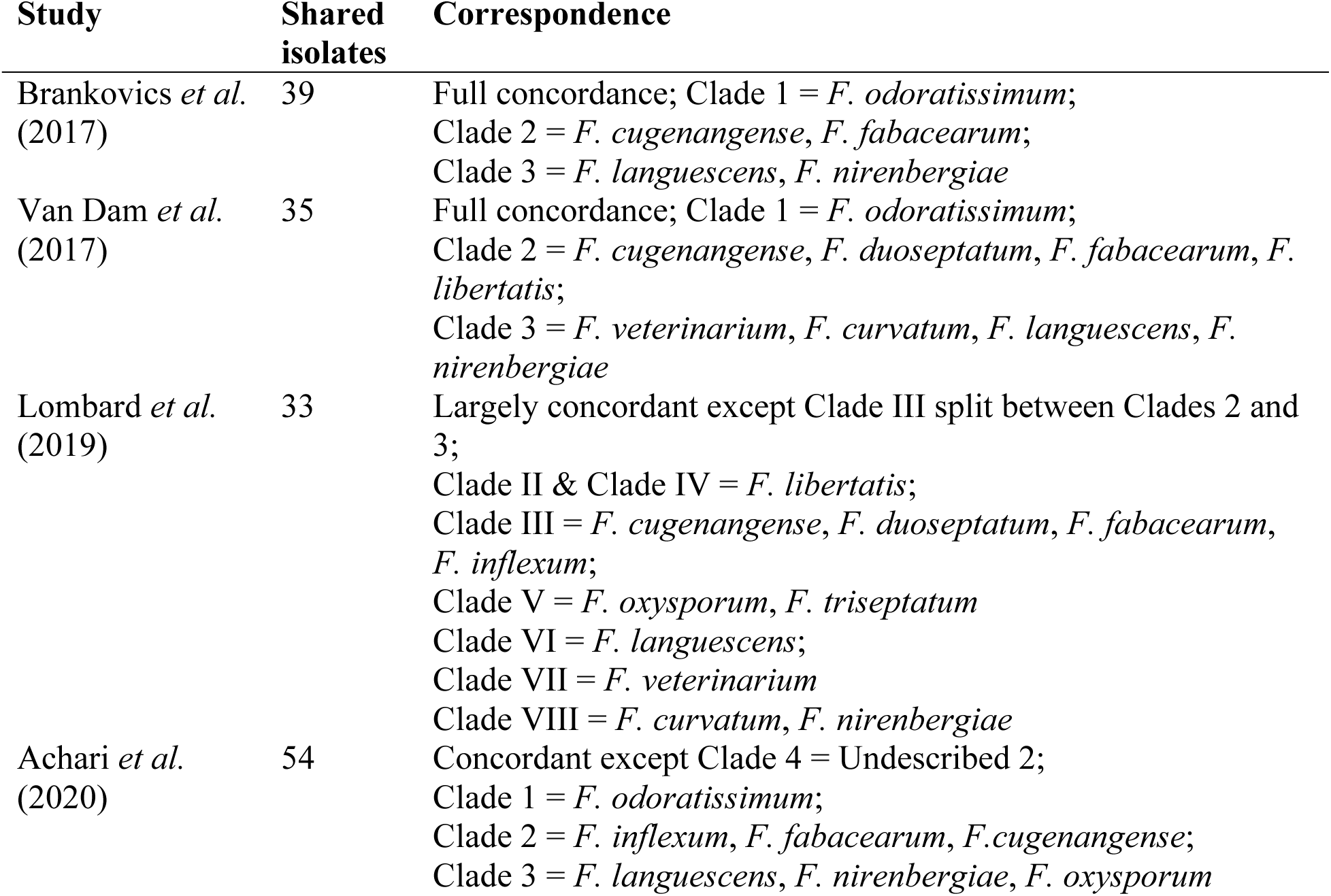
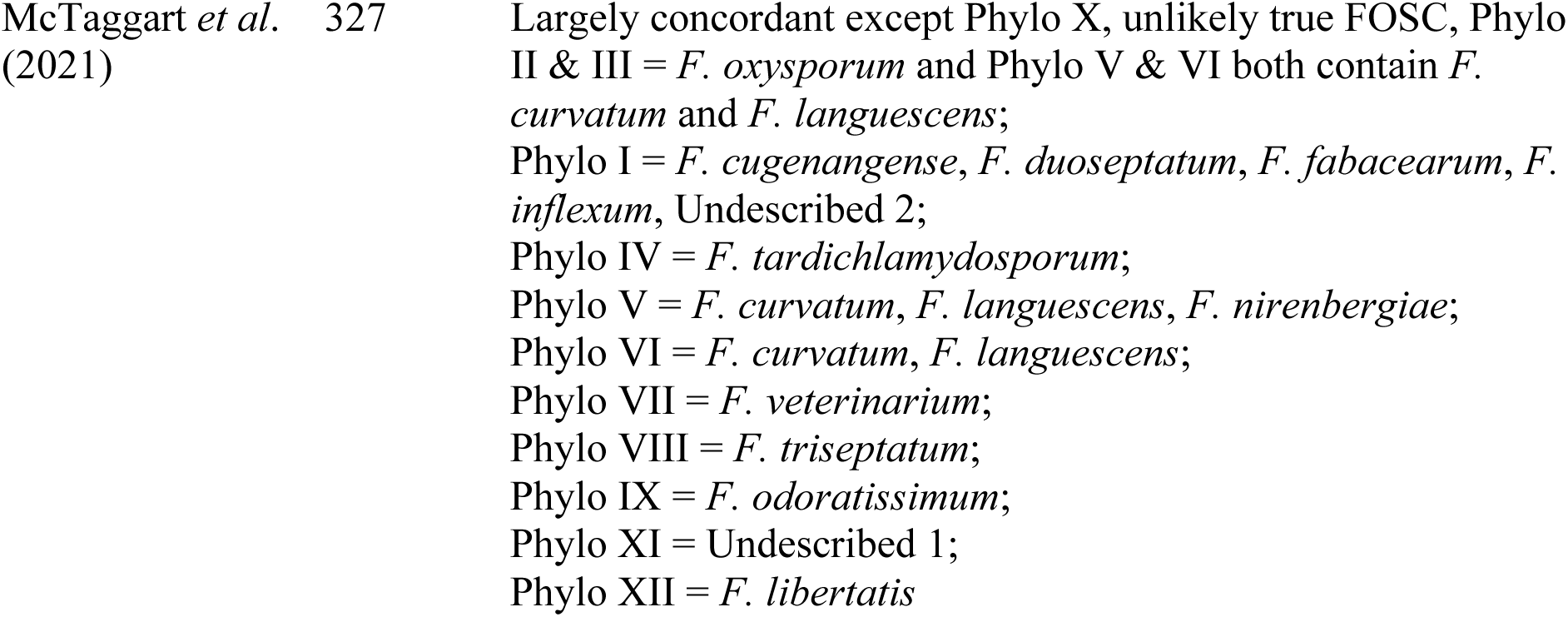
Putative species delineated in FOSC by our phylogenomic studies largely correspond to previously described taxa.

**Fig. 3.**
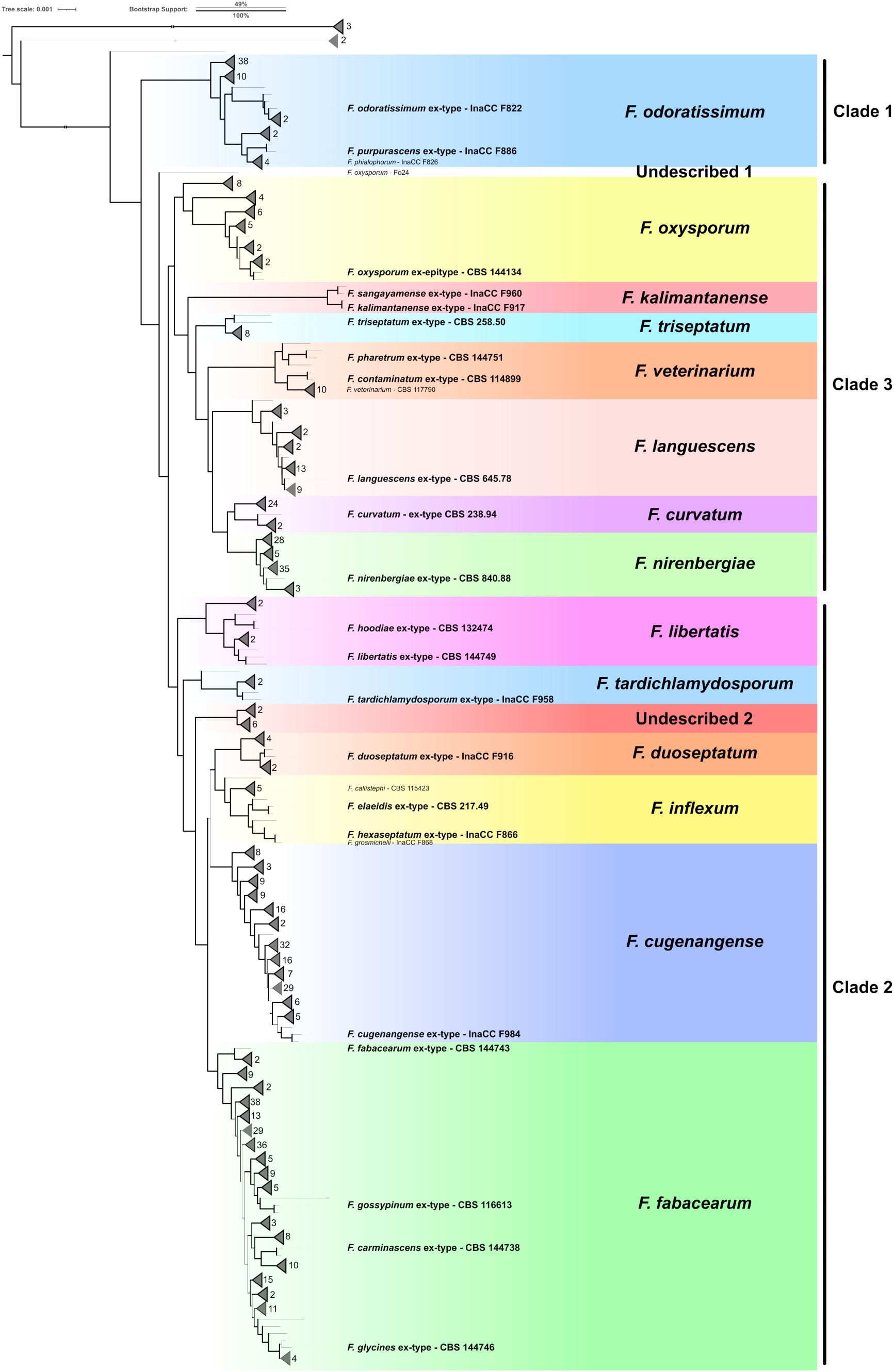
The *Fusarium oxysporum* species complex based on a phylogenomic analysis contains fourteen described species as well as two undescribed lineages. The phylogenetic tree was computed using a concatenation of 2493 BUSCO genes that are conserved in a single copy across all 663 included genome assemblies. Species are indicated with different shading, and bootstrap support is indicated with branch thickness. The branches leading to the outgroup isolates of *Fusarium redolens* and to the non-FOSC isolates have been shortened for visualization purposes. Clades without any type-isolates have been collapsed for visualization purposes, and the numbers next to triangles indicate the number of isolates within each collapsed clade. Representative isolates of described species without an ex-type in this analysis are also labelled. See Sup. Fig. 3 for the complete phylogenetic tree.

### Major lineages in the *Fusarium oxysporum* species complex are not driven by minority gene subsets

Although the maximum-likelihood analyses (Fig. 3) yielded high ultrafast bootstrap support for the 16 lineages, we sought to evaluate the underlying genomic consensus supporting these. We therefore quantified gene and site concordance factors (sCF and gCF), which ranged between 0.05 and 99.1 % and 0 to 100 %, respectively, indicating that some branches in the phylogeny were only supported by a minority of genes and parsimony informative sites. Notably, gCF and sCF for the 16 main lineages ranged from 2.25 to 87.9 and 30.6 to 85.9, respectively (Sup. Fig. 3). Thus, while these 16 lineages are well-supported by maximal ultrafast bootstrap values and had previously been recovered by independent analyses (Brankovics *et al*. 2017, van Dam *et al*. 2017, Lombard *et al*. 2019, Achari *et al*. 2020, McTaggart *et al*. 2021), their comparatively low gCF and sCF values suggest substantial underlying discordance within the marker set.

To further challenge the robustness of the inferred phylogeny and the species delineation, especially given the limited gCF and sCF support for some nodes, we derived a total of 38 phylogenetic trees based on 19 unique subsets of our 2,449 BUSCO markers, using both concatenated maximum-likelihood (IQ-TREE) and coalescence (ASTRAL) frameworks. These subsets were selected to evaluate the effect of markers with relatively low and high parsimony-informative content, as well as randomly sampled markers (Fig. 4A). In 36 of 38 (94.7 %) subset phylogenies, all 16 putative species were recovered as monophyletic clades (Fig. 4). Beyond species-level monophyly, we also assessed whether each species fell within its expected overarching clade relative to the maximum-likelihood and coalescence reference analyses; this was the case in 28 of 38 (73.7 %) subset trees, and for all species except *F.* o*xysporum* s. str. this was the case for 38/38 (100 %) of trees. Thus, *F. oxysporum* was the only species to shift clade placement, moving from Clade 3 to Clade 2 in ten subset trees, for example, in ordered subsets 4 and 7 under both the maximum-likelihood and coalescence approaches, and in random subsets 1, 7, and 8 under the coalescence approach (with only a subset of *F. oxysporum* isolates shifting in two of these cases, and the isolates split between Clades 2 and 3 in random subsets 5 and 6; Fig. 4C). Taken together, these results indicate that our putative species are robustly recovered as monophyletic lineages across the various data subsets, with only minor and localized perturbation in the higher-level clade placement of *F. oxysporum*.

**Fig. 4.**
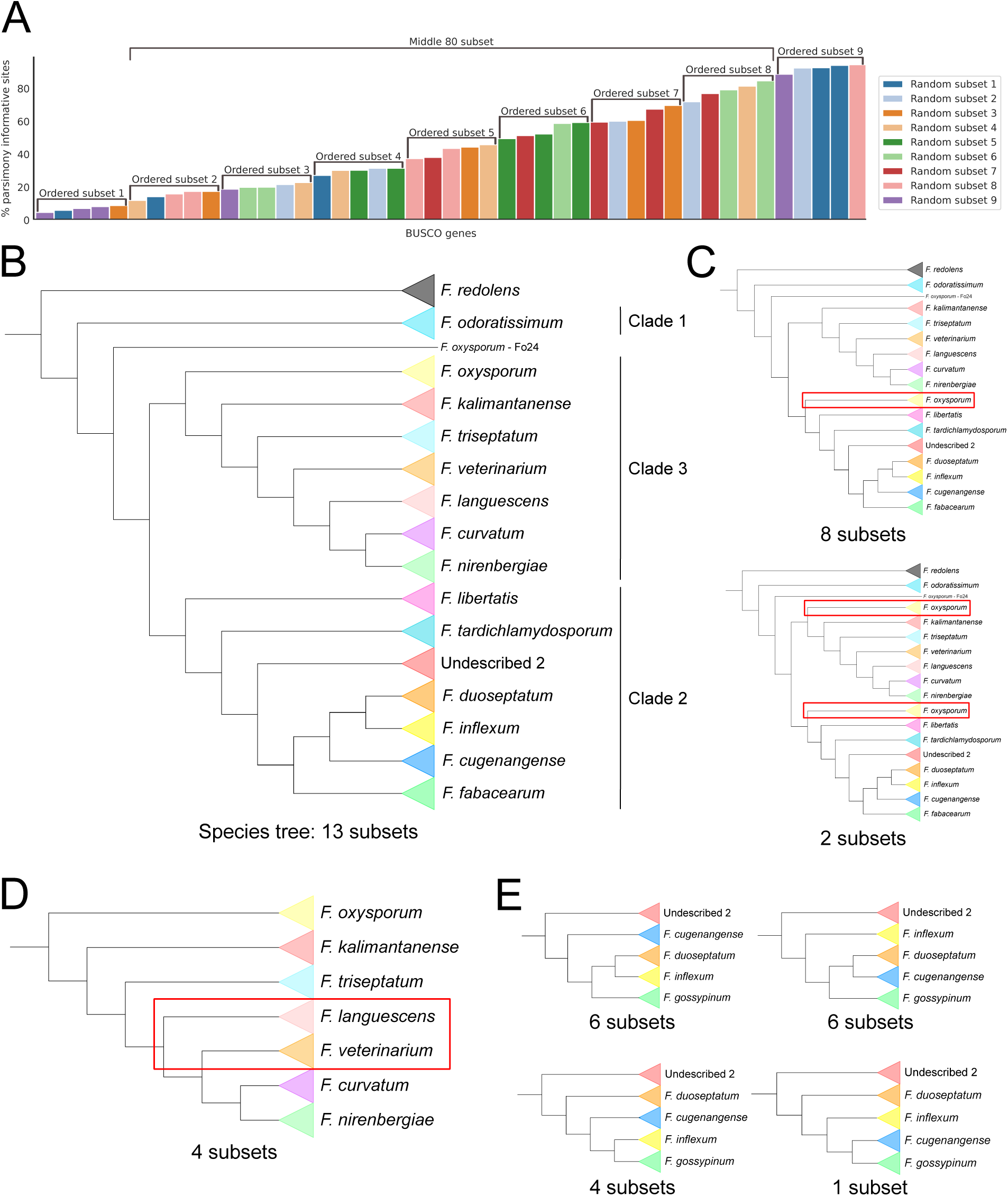
FOSC species are recovered as monophyletic clades in nearly all phylogenetic analyses. **A.** Graphical representation of the 19 different subsets created from the BUSCO gene dataset. Each subset was subject to both maximum-likelihood- and coalescence-based phylogenetic analysis, yielding 38 phylogenies trees derived from a subset of markers. **B.** Summary of the species phylogeny (Fig. 3), indicating that 13/38 (34.2 %) subset phylogenies were identical to the species tree. **C–E.** Graphical overview of the observed discrepancies between the species tree (Panel B) and phylogenetic analyses resulting from subsets of the markers. Multiple discrepancies relative to the species tree can be observed simultaneously within a single subset tree. **C.** Two observed movements of *F. oxysporum* completely or partially moving from clade 3 to clade 2. **D.** An observed shift of *F. languescens* and *F. veterinarium*. **E.** Four observed alternative topologies for species *F. duoseptatum, F. inflexum, F. cugenangense,* and *F. fabacearum*.

### FOSC lineages have become clonal after descent from a sexual ancestor

To assess the degree of recombination within the FOSC and to determine whether the low support (gCF and sCF) values (Sup. Fig. 3) might reflect complex, reticulate gene flow, we subjected the data to a SplitsTree analysis (Huson & Bryant 2024) (Fig. 5A). The resulting network revealed 16 separated clusters with minimal reticulation. Reticulation was concentrated primarily near the root of the network and within Clade 3, suggesting ancient recombination in the ancestral FOSC that has since become (largely) clonal. This interpretation is supported by evidence of sexual recombination in many other *Fusarium* species complexes, including *N. solani* (= *F. solani)*, *F. fujikuroi,* and *F. graminearum,* suggesting that the ancestor of the FOSC was likely a sexually recombining species, and the contemporary FOSC has (largely) lost that ability (Covert *et al*. 2007, Talas & McDonald 2015, Fourie *et al*. 2018). This is further suggested by the repurposing of mat loci within the FOSC (Vitale *et al*. 2026). Notably, the network topology, with its 16 clusters, closely matches the topology of the maximum-likelihood-based tree topology (Fig. 3), suggesting that low gCF and sCF values likely reflect stochastic gene-tree error caused by for example incomplete lineage sorting rather than genuine evolutionary conflict, further corroborating the here delineated putative species within the FOSC.

**Fig. 5.**
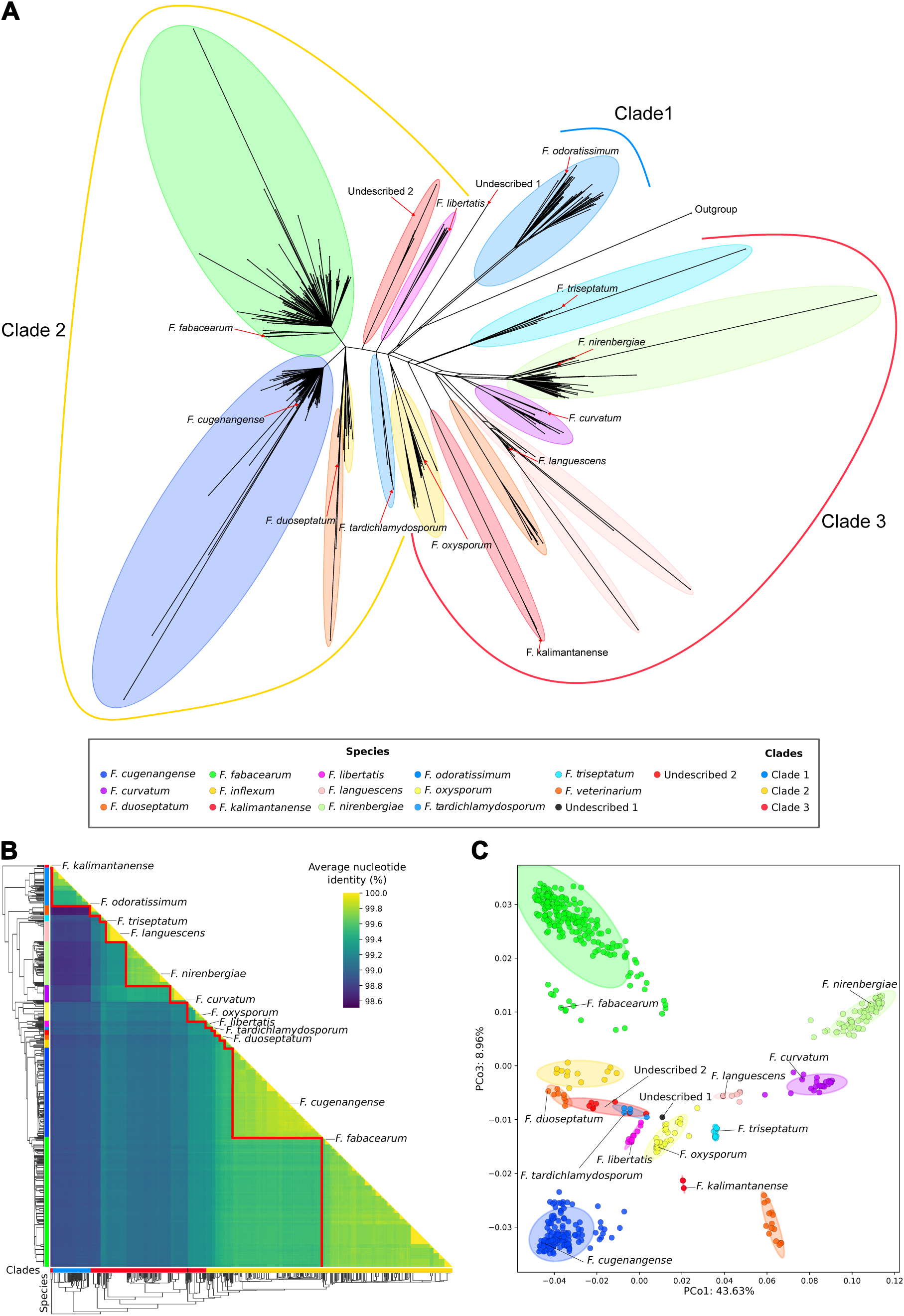
Putative species within the FOSC are also recovered using whole-genome comparisons. **A.** SplitsTree network, based on 2493 BUSCO genes, suggests minimal signal for reticulation, e.g., through recombination or introgression; netting or parallel lines indicate reticulation, while single lines represent clonality. The outgroup branch was manually shortened and collapsed for improved visualization. Putative species (Fig. 3) were mapped onto isolates for visualization subsequent to the analyses, and arrows/lines indicate type isolates for putative species if available (applies for A-C). **B.** Whole-genome nucleotide identity analysis recapitulates previously identified species (Fig. 3). Nucleotide-identity matrix, obtained from genome-wide alignments of core genomic regions of all 597 FOSC isolates in our dataset, were reordered using average linkage clustering over genome dissimilarity. **C.** Principal Coordinates Analysis (PCoA) based on biallelic SNPs consistently recovers previously identified species (Fig. 3) in Clade 2 and Clade 3. For clarity, we excluded Clade 1 (*F. odoratissimum*) as it strongly separates from Clade 2 and 3, limiting differentiation within Clade 2 and 3 (see Sup. Fig. 4A for PCoA of all isolates). We here showed PCo3 instead of PCo2 as this improved visual differentiation of putative species; difference in variance explained was only 0. 92 % (PCoA of PCo1 and PCo2 shown in Sup Fig. 4B).

Our analysis found little evidence of recent or ongoing recombination within the FOSC (Fig. 5A), contrasting prior reports from core and mitochondrial genome patterns (Brankovics *et al*. 2020, McTaggart *et al*. 2021, Fayyaz *et al*. 2023). This discrepancy may partly reflect differences in methodology. McTaggart *et al*. (2021) performed SplitsTree on only 40 genes, which is potentially more susceptible to variation at single loci [e.g., incomplete lineage sorting (ILS)], whereas we used a concatenated alignment of 2493 BUSCO genes. However, the strong conservation of BUSCO genes within the *Nectriaceae* may itself limit the detection of recombination, especially if recombination is restricted to small, closely related subgroups of isolates, as proposed by McTaggart *et al*. (2021) as well as Fayyaz *et al*. (2023). If so, BUSCO genes might be too conserved to yield detectable gene-tree incongruence. Still, ongoing (para)sex among closely related isolates would fall within our proposed species boundaries and would thus not conflict our species delineations.

### Nucleotide identity is higher within than between putative FOSC species

To incorporate information from the whole genome, rather than solely focusing on conserved BUSCO genes, we computed pairwise nucleotide identities between genome assemblies of 657 FOSC isolates in our dataset, excluding the *F. redolens* outgroup isolates as well as the three isolates we initially identified as likely non-FOSC (GCA_014324465.1, GCA_019157295.1, and GCA_024865645.1). To exclude any potentially conflicting genomic signal from accessory chromosomes, which are known to be able to transfer horizontally between FOSC isolates (Ma *et al*. 2010), we first removed accessory chromosomes and regions from each genome assembly; the core genome ranged between 36.5 and 50.2 Mb, with an average of 41.5 Mb. The average core genome size covered exactly once across all pair-wise genome alignments was 21.8Mb, corresponding to approximately 52.7 % of the average core-genome. Absence of alignments could result either from repetitive elements, which are likely to be mapped multiple times, or from extensive divergence between isolates. To minimize the risk of spurious signal from non-homologous or paralogous mappings, we applied a strict identity threshold of max 0.1 % sequence divergence, resulting in just over half of the core genome being mapped between any given pair of genomes. Across the mapped genomic regions, the mean number of SNPs found between two isolates was 151,769 (range: 0 - 298,896). Notably, no SNPs were only recovered in ten out of 430,992 comparisons, and these occurred exclusively between two isolates of the same putative species (six comparisons within *F. oxysporum* and four within *F. languescens*). The average nucleotide identity among isolates assigned to the same putative species in the species tree was 99.74 % (median: 99.73 %) whereas the average nucleotide identity between isolates of different putative species was notably lower at 99.16 % (median 99.11 %). Clustering based on nucleotide identity, and subsequent overlaying of the here delineated putative species (Fig. 3) onto the resulting dendrogram, recovered all 16 putative species as distinct clusters (Fig. 5B). Clades 1 and 2 were likewise fully recovered in the clustering, while Clade 3 was not recovered as a coherent group since *F. oxysporum* clustered closer to Clade 2 than to the rest of Clade 3, mirroring the pattern observed for a few of the subset trees, and *F. kalimantanense* did not group with the rest of the clade (Fig. 5B). Thus, the 16 previously delineated lineages could also be successfully recovered using whole-genome alignments.

To further corroborate our species delineation, we obtained single nucleotide polymorphisms (SNPs) from whole (core) genome alignments of all 657 FOSC assemblies against the core genome of the reference isolate *F. oxysporum* f.sp. *lycopersici Fo*4287, yielding in total 2,148,320 unique, biallelic SNPs unique relative to the *Fo*4287 core genome; 957,294 (44.5 %) were removed after filtering for a minimum allele frequency of at least 0.5 %, leaving 1,191,026 SNPs for downstream analysis. We then used this filtered data for Principal Coordinate Analysis (PCoA). PCo1 captured 34.22 % of the total variation, PCo2 captured 20.43 %, and PCo3 captured 7.24 %. PCo2 separates Clade 1 from the other two clades (Sup. Fig. 4A). Since Clade 1 currently contains only a single putative species, we repeated the analysis excluding Clade 1 to further emphasize the separation within Clade 2 and 3. After removing Clade 1, our dataset contained 1,034,087 SNPs, and PCo1 captured 43.63 % of variation, PCo2 captured 9.61 %, and PCo3 captured 8.69 % (Fig. 5C). Given the relatively small difference in variance captured between PCo2 and PCo3, both axes were visualized alongside PCo1 (Fig. 5C and Sup. Fig. 4B). Species within Clades 2 and 3 were most clearly separated by the combination PCo1 and PCo3, where PCo1 primarily distinguished between clades and PCo3 differentiated most notably between *F. fabacearum*, *F. nirenbergiae*, *F. cugenangense*, and *F. veterinarium* (Fig. 5C). Notably, *F. oxysporum* clustered between Clades 2 and 3 on PCo1. Using two standard deviations from the centroid of species as a proxy of the diversify within a species of (shown as an ellipse in the PCoA), only ‘Undescribed 2’ partially overlapped with *F. duoseptatum* and with *F. tardichlamydosporum*. In contrast, no other isolate was positioned inside these intervals of another species, and all ex-type isolates consistently clustered within their respective species, strongly suggesting that the putative species delineated previously (Fig. 3) are indeed robust and can be recovered using a variety of whole-genome analyses.

### Marker genes traditionally used to differentiate species in FOSC do not contain sufficient information for accurate species identification

The genetic markers *tef1*, *rpb2*, *rpb1*, *CAM*, *tub1*, and ITS are commonly used to identify fungal isolates up to species level and represent the most frequently sequenced marker genes for FOSC isolates (Table 1), and many phylogenetic studies and assignments of novel isolates to existing or even novel species are based solely on (subsets of) these markers (O’Donnell *et al*. 1998, 2004, Baayen *et al*. 2000, Laurence *et al*. 2011, Lombard *et al*. 2019, Achari *et al*. 2020, Nephalela-Mavhunga *et al*. 2021, Zhang *et al*. 2023, Tan & Conroy 2024). To determine the power of these genes to recover the here defined putative species within FOSC, we extracted their sequences from the 663 whole-genome assemblies and conducted phylogenetic analyses of the concatenated markers. In a maximum-likelihood phylogenetic analysis, only 12 out of 16 putative species were recovered as monophyletic clades (Fig. 6B). Moreover, even the well-established separation between Clades 2 and 3 were not correctly recaptured, as isolates of *F. libertatis* and *F. oxysporum* clustered outside of Clades 2 and 3, respectively, with both putative species split into three separate clusters distributed across the phylogeny. Similarly, *F. tardichlamydosporum* was split into two distinct lineages, and *F. fabacearum* was split into three distinct clusters. The latter was caused by the insertion of both *F. inflexum* and *F. duoseptatum* within this putative species relative, breaking up *F. fabacearum* as a monophyletic species. Thus, the six common marker genes were insufficient for accurate and reproducible identification of the 16 putative delineated species and thus should only be used cautiously for species delineation in FOSC.

### Only five additional BUSCO-gene markers alongside *tef1* and *rpb2* are necessary for species-identification of FOSC isolates

A simple and accurate species identification is indispensable for any taxon, and particularly for phytopathogenic fungi as impactful as those found in the FOSC (Hariharan & Prasannath 2021). We thus sought to identify BUSCO genes as additional diagnostic markers for species delineation in FOSC. We first selected *tef1* and *rpb2* as the starting point because they contain the most parsimony informative positions among the six commonly used marker genes and are also the most frequently sequenced (Table 1). We then supplemented a phylogeny based solely on *tef1* and *rpb2* with additional BUSCO marker genes and identified a minimal set of five additional BUSCO genes that, when combined with *tef1* and *rpb2*, together contained at least one parsimony informative site per putative species and consequently fully recaptured both the three canonical clades and all 16 putative species delineated here (Fig. 6C). This alternative marker set contained a total of 1434 parsimony-informative positions across a combined alignment length of 12,164 bases (11.7 % parsimony informative sites), including *tef1* and *rpb2* (Sup. Table 5). This suggests that BUSCO genes provide a useful resource to augment traditional marker genes with additional sequences to support species identification, without a significant increase in methodological complexity. Future studies should aim to develop these genes into PCR-amplifiable markers which can be sequenced for phylogenetic analyses where whole genome sequencing is not possible. In addition, these markers should also be tested for usefulness in other *Fusarium* species complexes.

**Fig. 6.**
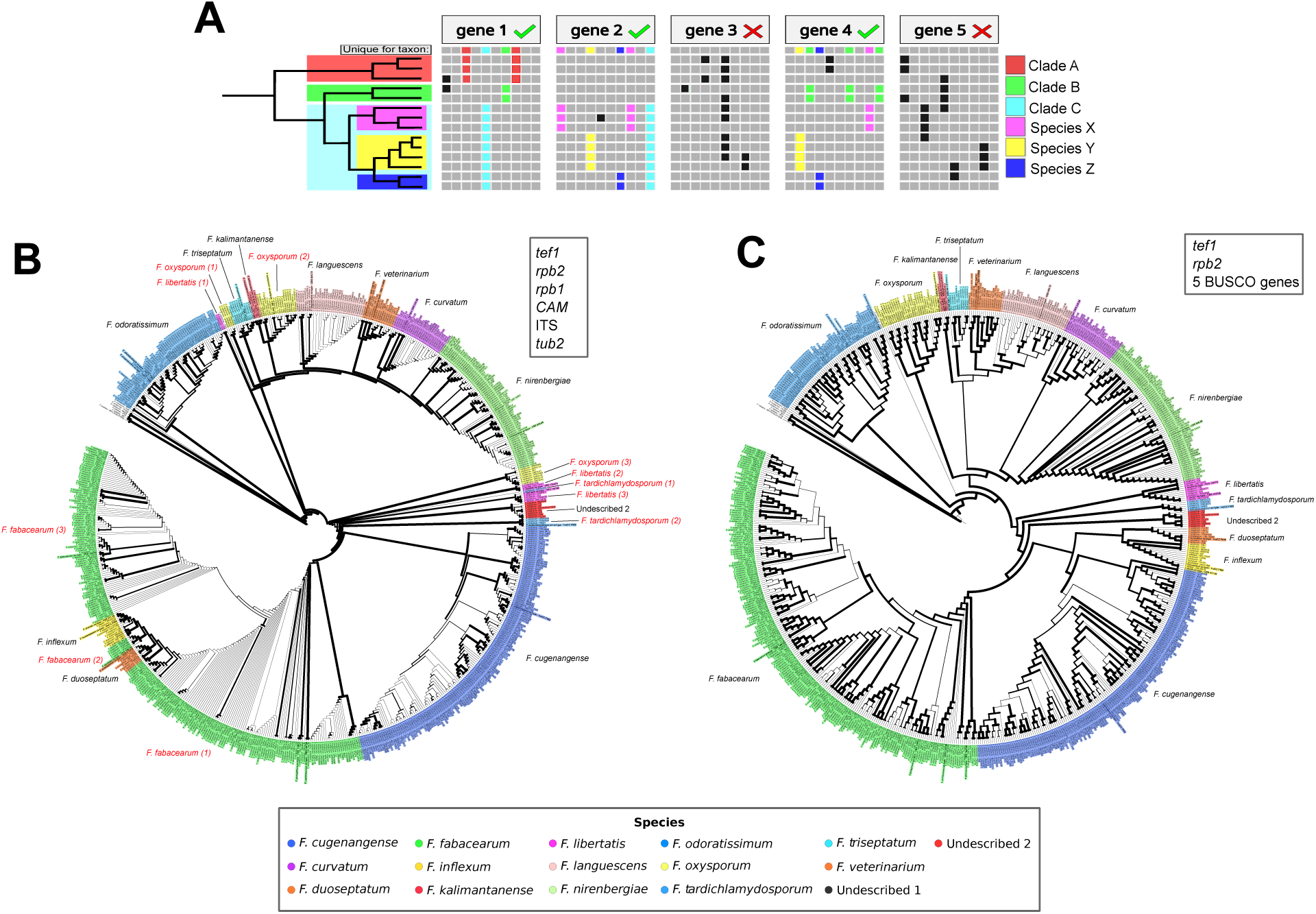
BUSCO markers are better suited for species identification in FOSC compared with traditional markers. **A.** Schematic overview of selection process of potential BUSCO marker genes suitable to differentiate the here delineates species and lineages. Genes were scanned per alignment position for residues conserved within a species but different in all other isolates. A minimal combination of genes was then identified with informative residues for all putative species. **B.** Maximum-likelihood phylogeny of 663 FOSC isolate dataset based on six common marker genes: *tef1*, *rpb2*, *rpb1*, *CAM*, ITS, and *tub2*. Putative species identities as defined in Fig. 3 are shown as the coloured background labels; putative species names are placed next to the respective lineages. Black names are recovered as monophyletic clades, while red names indicate species not recovered as such. Numbers between brackets after putative species names indicate different subgroups of respective species. Branch thickness indicates bootstrap support. **C.** Maximum-likelihood phylogeny based on *tef1*, *rpb2*, and five BUSCO marker genes (50269at110618, 100592 at110618, 121884 at110618, 30696 at110618 and 119914 at110618). Branch thickness indicates bootstrap support. Names and label backgrounds indicate putative species, as defined in Fig. 3.

## CONCLUSIONS

The taxonomy of the *Fusarium oxysporum* species complex (FOSC) has become increasingly blurred, confusing and difficult to apply, largely due to the limited resolving power of existing marker genes and conflicting, difficult to compare results generated by previous studies (O’Donnell *et al*. 1998, 2004, Baayen *et al*. 2000, Laurence *et al*. 2011, 2014, Brankovics *et al*. 2017, van Dam *et al*. 2017, Lombard *et al*. 2019, Achari *et al*. 2020, McTaggart *et al*. 2021). Stabilization of FOSC taxonomy is a pressing priority, given that the complex is among the most impactful and threatening groups of phytopathogens worldwide (Dean *et al*. 2012, Bhunjun *et al*. 2024). Here, we leveraged public whole-genome sequencing (WGS) data, complemented by novel genome assemblies of ex-type isolates, to propose a taxonomic reorganization of FOSC aimed at clarifying species boundaries within the complex. Using 2,493 markers, we resolved 16 lineages, leading to the synonymy of 12 species, distributed across three well-described clades (O’Donnell *et al*. 1998, 2004, Baayen *et al*. 2000, Brankovics *et al*. 2017, van Dam *et al*. 2017, McTaggart *et al*. 2021). To further corroborate the robustness of this species delineation, we evaluated our phylogenetic model using measures of introgression, genome-wide nucleotide identity, and genome-wide SNP data, all of which suggest that these lineages are robust taxonomic units. Finally, we identified novel marker genes capable of assigning newly sampled strains to these established lineages, providing a practical foundation for the future description of additional species. We thus anticipate that this revised phylogeny will offer a useful framework for the community and help resolve the existing confusion surrounding FOSC taxonomy.

Our study is, to our knowledge, the first to not only focus on a few strains or employ limited phylogenetic methods, but to involve whole genomes of a large and diverse collection of FOSC isolates, including more than 20 ex-type isolates. This allows us now to start consolidating FOSC diversity into a single, unified phylogenetic framework. Our analyses are largely consistent with previous work as we recapitulate clades as well as lineages described in current literature (Brankovics *et al*. 2017, van Dam *et al*. 2017, Lombard *et al*. 2019, McTaggart *et al*. 2021). The here accepted 16 species within FOSC have been described and/or recaptured as monophyletic taxa independently in previous studies, emphasizing the need for large-scale phylogenomic approaches, as similarly recently employed in *Aspergillus* (Steenwyk *et al*. 2019) and *Colletotrichum* (Ma *et al*. 2025). Although each species is well supported as an independent taxon, relationships between some species remain difficult to resolve. Placement of *F. oxysporum* relative to the other species proved especially challenging, as did resolving relationships between *F. languescens*/*F. veterinarium* and *F. curvatum*/*F. nirenbergiae*, and among *F. duoseptatum*, *F. inflexum*, *F. cugenangense*, and *F. fabacearum*. In each case, the lineages appear to have diverged in rapid succession, consistent with the limited observed variation among them. Such rapid divergence is known to hinder species delimitation (Struck *et al*. 2018), primarily through incomplete lineage sorting (ILS), which likely explains the discordance among subset trees and the lower support at these nodes (Degnan & Rosenberg 2009, Liu *et al*. 2015, Weber *et al*. 2019). The ambiguous placement of *F. oxysporum* may carry particular taxonomic significance. We retained it within Clade 3, consistent with current literature that places this species with other species in Clade 3 (Lombard *et al*. 2019, Achari *et al*. 2020). However, we observed variable placement across analyses as well as greater pairwise nucleotide identity to Clade 2 isolates, which warrants further investigation. A more densely sampled FOSC backbone, spanning a wider range of lifestyles and geographic origins, may ultimately place *F. oxysporum* within Clade 2 or even split it, warranting the recognition of two additional species. Yet, as most of our analyses support *F. oxysporum* as a monophyletic entity and no additional species have to our knowledge been described within this species, we retain *F. oxysporum* as a single species.

Across our analyses, several long attracting branches appear in both the phylogenetic trees and the SplitsTree analysis, which can affect the results of comparative genomic analyses (Szánthó *et al*. 2023). Such branches could stem from poor alignment or low sequencing quality. However, we also observed these branches in Figure 1, where we manually verified the alignment, reducing the likelihood of a technical artifact. Alternatively, the distance between strains may be exaggerated simply because more closely related strains have not yet been sequenced. Moreover, since a substantial number of FOSC strains are endophytic or saprophytic, considerable biodiversity may be missing from our sampling, making these branches appear disproportionately long.

Our phylogenomic analyses of the FOSC carry important implications not only for taxonomy but also related fields. We provide a robust, stable framework for species identification, grounded in the available diversity of whole-genome assemblies and complemented by newly sequenced ex-type isolates, resolving the inconsistent and conflicting species definitions that have hindered the field, further supporting direct comparison across different taxonomic studies. Within this framework, we synonymize 12 species, substantially simplifying FOSC taxonomy. We show that substantial genomic diversity exists within the FOSC, and that even within a single putative species, commonly used marker genes vary considerably, rendering these insufficient for reliable species recognition or novel species description. Our framework instead offers a platform to guide the description of novel species, in contrast to current largely ad hoc practice, where species are often delineated using minimal divergence at a few poorly resolved marker genes within a sparsely sampled tree (Zhang *et al*. 2023, Tan & Conroy 2024). We further show that phylogenomic data can identify improved markers. By supplementing *tef1* and *rpb2* with as few as five additional BUSCO genes, we fully recovered the 16 species delineated here. Pending experimental validation through primer design, these markers may also enable rapid, cost-effective molecular identification of FOSC isolates. Thus, our analysis fundamentally changes the general approach to FOSC taxonomy, moving from poorly resolving marker genes to the incorporation of whole-genome data and, potentially, novel, targeted marker genes.

Building on this framework, we see two clear priorities for the community going forward: First, only 22 of the 51 currently accepted FOSC species (43 %, per fusarium.org) are represented by ex-type isolates in our study, all of which were newly sequenced here. To our knowledge, no genomes from (ex-)type material of FOSC species were available in public repositories at the time of writing. It is thus essential that the remaining ex-type isolates will be sequenced, either via whole-genome sequencing or based on the identified minimal BUSCO markers, such that future studies will be able to place remaining species within this framework, resolve names, and synonymize taxa where appropriate. We further recommend that novel species descriptions rely on whole-genome analyses within well-sampled phylogenies, and that whole-genome assemblies become the standard for taxonomic practice as they grow more commonly accessible, serving as a template for species delineation in other fungal lineages more broadly. Second, our dataset, and FOSC research as a whole, is currently weighted toward putatively phytopathogenic isolates (311/663 (46.9 %) isolates in our dataset, likely an underestimate (given often incomplete annotation), while growing evidence suggests that most FOSC isolates are in fact non-pathogenic, adopting saprotrophic or rhizosphere-associated lifestyles (Edel-Hermann & Lecomte 2019, Martínez-Soto *et al*. 2023). Expanding sampling to better represent this non-pathogenic diversity will be key for future work, as it would broaden the phylogenetic backbone underlying species delineations and help disentangle lifestyle-associated biases. For example, it is conceivable that pathogenic lineages could be disproportionately often clonal, having arisen from less-frequently sampled recombining populations. Expanded sampling may thus also help clarify the limited recombination signal we observed here, despite previous proposals of (para)sexuality in FOSC (Brankovics *et al*. 2017, McTaggart *et al*. 2021, Fayyaz *et al*. 2023). We therefore encourage future sampling efforts to prioritize lifestyle diversity alongside phylogenetic breadth, and to incorporate these isolates into the taxonomic framework as it continues to develop. Doing so has strong potential to refine our understanding of speciation and species boundaries within the FOSC and will open novel avenues of research.

## Supporting information

Supplemental Tables (combined)

## Citation

Bell J, van Os J, Thines M, Costa MM, Collemare J, Groenewald JZ, Crous PW, Seidl MF (2026). Elucidating cryptic species in the *Fusarium oxysporum* species complex. *Studies in Mycology* xxxx: xx–xxx. doi: xxxxxx

## Corresponding editor

F. Liu

## DECLARATION ON CONFLICT OF INTEREST

The authors include members of the Editorial Board of Studies in Mycology. They were not involved in the journal’s review of, or decisions related to, this manuscript.

## SUPPLEMENTARY DATA DESSCRIPTIONS

**Fig. S1.** A phylogenetic tree based on six commonly used marker sequences is insufficient to robustly delineate species in the *Fusarium oxysporum* species complex. The maximum-likelihood based phylogenetic tree was inferred using a concatenated nucleotide alignment of *tef1, rpb2, rpb1, CAM*, ITS, and *tub2* obtained from 287 FOSC isolates and outgroup isolate *Fusicolla matuoi* CBS 581.78 from fusarium.org. Type isolates are highlighted in bold, branch thickness indicates bootstrap support, and the outgroup branch was shortened for visualization purposes. The outer ribbon indicates isolates with a clade annotation from previous studies, from which monophyletic species were extrapolated, as emphasized by the background shading. **Left:** Branch length represents substitution per sites and isolates selected for whole-genome sequencing (Fig. 2) are emphasized with a red star.

**Fig. S2.** Individual phylogenetic trees of commonly used marker sequences contrast current species delineations. Maximum-likelihood based phylogenetic trees were inferred from a nucleotide alignment per commonly used marker sequence (*tef1, rpb2, rpb1, CAM*, ITS, and *tub2*) obtained from Fusarium.org for up to 287 and outgroup isolate *Fusicolla matuoi* CBS 581.78, depending on data availability. Type isolates are highlighted in bold, branch thickness indicates bootstrap support, and the outgroup branch was shortened for visualization purposes. The outer ribbon indicates isolates with a clade annotation from previous studies

**Fig. S3.** The *Fusarium oxysporum* species complex based on a phylogenomic analysis contains 14 described species as well as two unnamed lineages. The phylogenetic tree was computed using a concatenation of 2493 BUSCO genes that are conserved in a single copy across all 663 included genome assemblies. Species are indicated with different shading, and bootstrap support is indicated on each branch following Ultrafast Bootstrap / gene Concordance Factor / Site Concordance Factor. The branches leading to the outgroup isolates of *Fusarium redolens* and to the non-FOSC isolates have been shortened for visualization purposes.

**Fig. S4.** Principal Coordinates Analysis (PCoA) based on biallelic SNPs are biased towards differentiating a subset of species. Points are coloured based on their putative species in Fig 3. If available, ex-types and Undescribed species are indicated with labels for putative species delineated in this study. **A)** PCoA of the complete dataset including all clades. PCo2 is strongly loaded to separate Clade 1 (*F. Odoratissimum*) from Clades 2 and 3. **B)** PCoA of Clades 2 and 3. PCo1 is s strongly loaded to separate Clades 2 and 3.

**Tab. S1.** Assembly statistics of 48 newly generated whole genome assemblies of FOSC isolates generated in this study, including 22 ex-type isolates.

**Tab. S2.** NCBI accession information, assembly statistics and putative species and clade definitions of all 663 Whole genome assemblies used in this study.

**Tab. S3.** Reference marker gene sequences used for BLAST searches for marker genes on whole genome assemblies.

**Tab. S4.** Parsimony informative sites statistics per BUSCO gene multiple sequence alignment.

**Tab. S5.** Descriptions of identified minimal combination of alternative marker genes sufficient for complete recovery of FOSC species tree.

## Notes

### Competing Interest Statement

The authors have declared no competing interest.

