## Supplementary material for "Consolidating species boundaries in the *Fusarium oxysporum* species complex based on genomic evidence": Figure S1

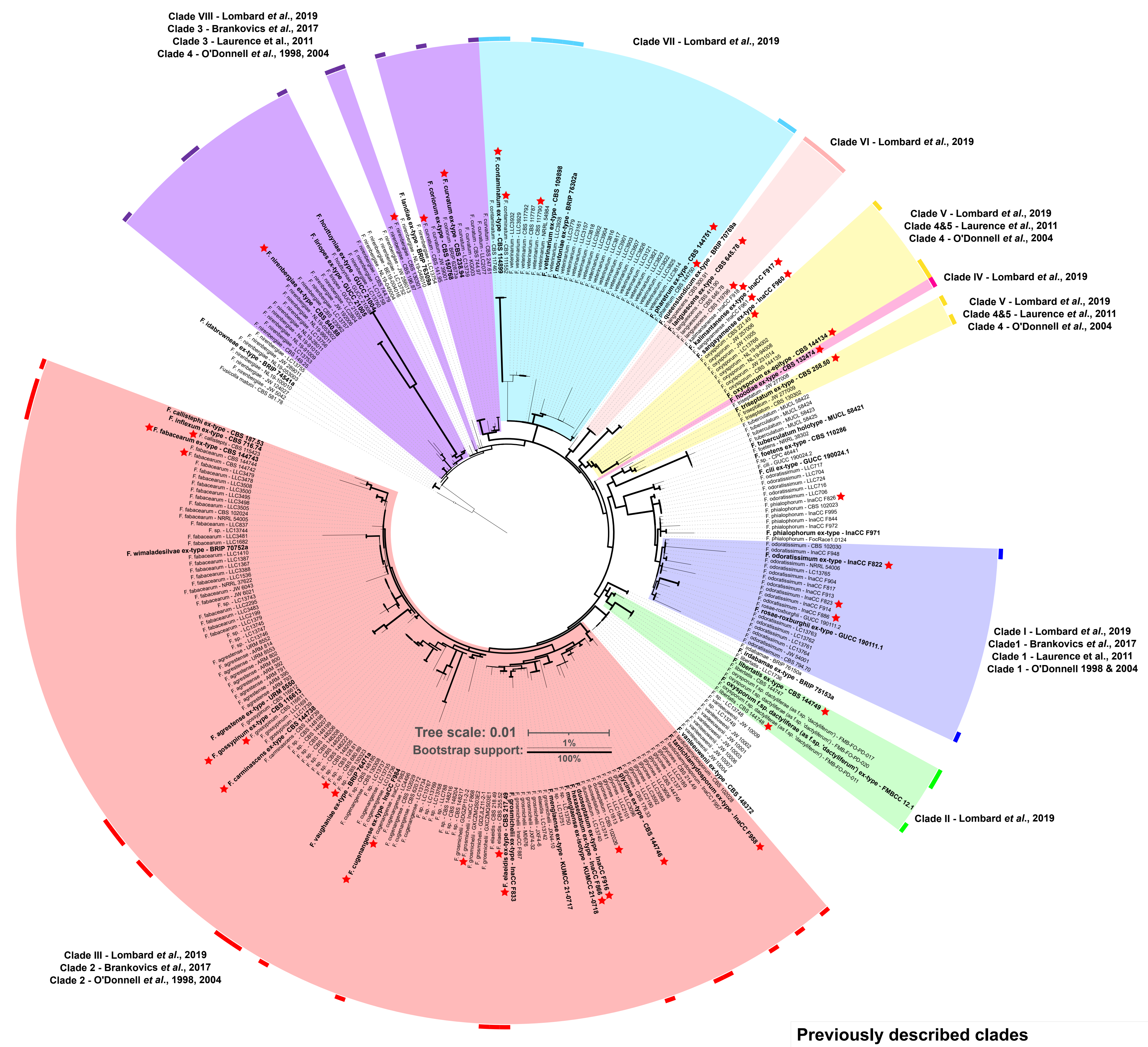


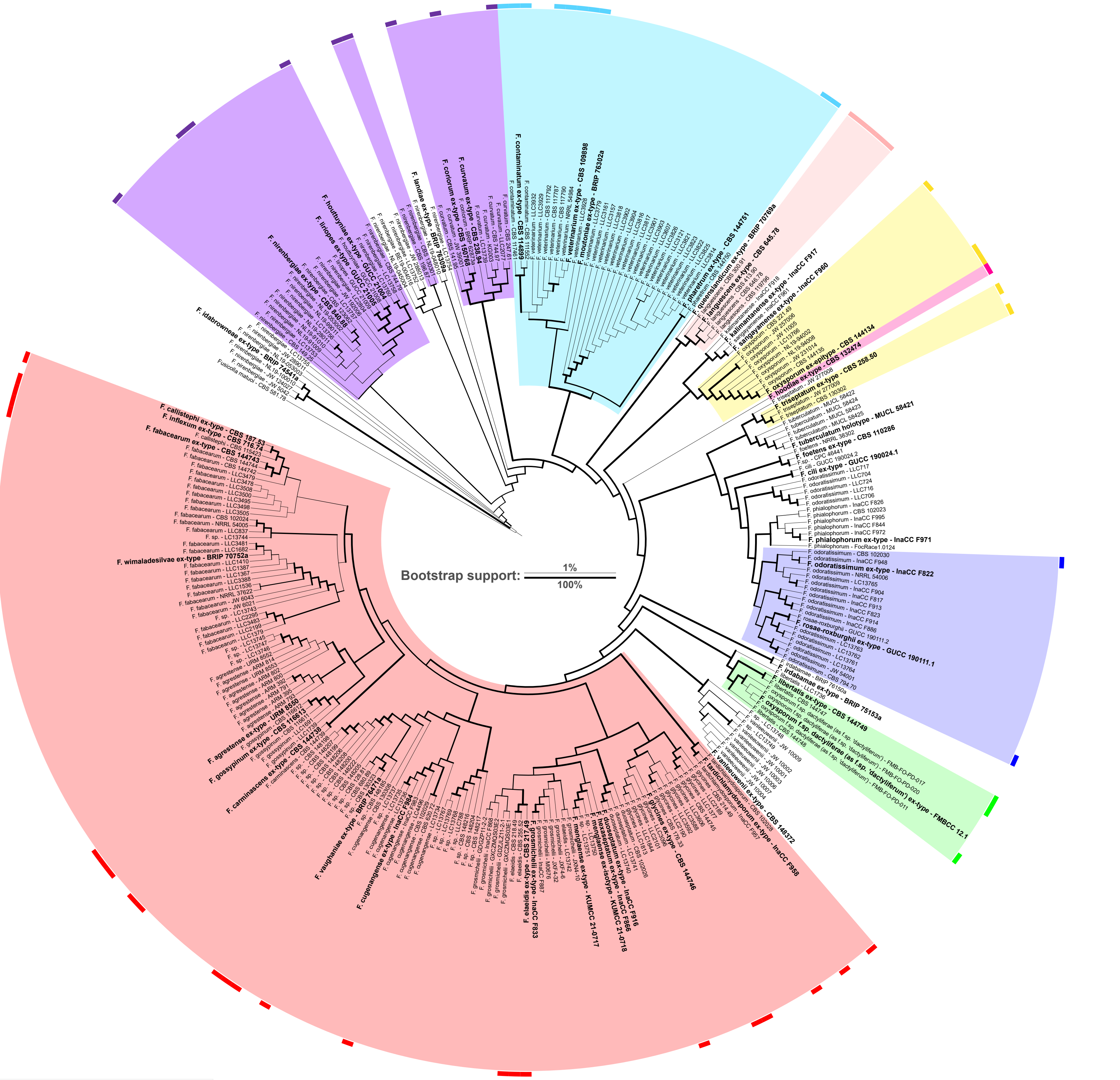


**Previously described clades**

| | Lombard et al., 2019 | O'Donnell et al., 1998 & 2004 | Laurance et al., 2011 | Brankovics et al., 2017 |
|---|---|---|---|---|
| | I | 1 | 1 | 1 |
| | II | | | |
| | III | 2 | | 2 |
| | IV | | | |
| | V | 4 | 4 & 5 | |
| | VI | | | |
| | VII | | | |
| | VIII | 4 | 3 | 3 |
