## Supplementary material for "Consolidating species boundaries in the *Fusarium oxysporum* species complex based on genomic evidence": Figure S2

### Calmodulin (*CAM* / *cmdA*)

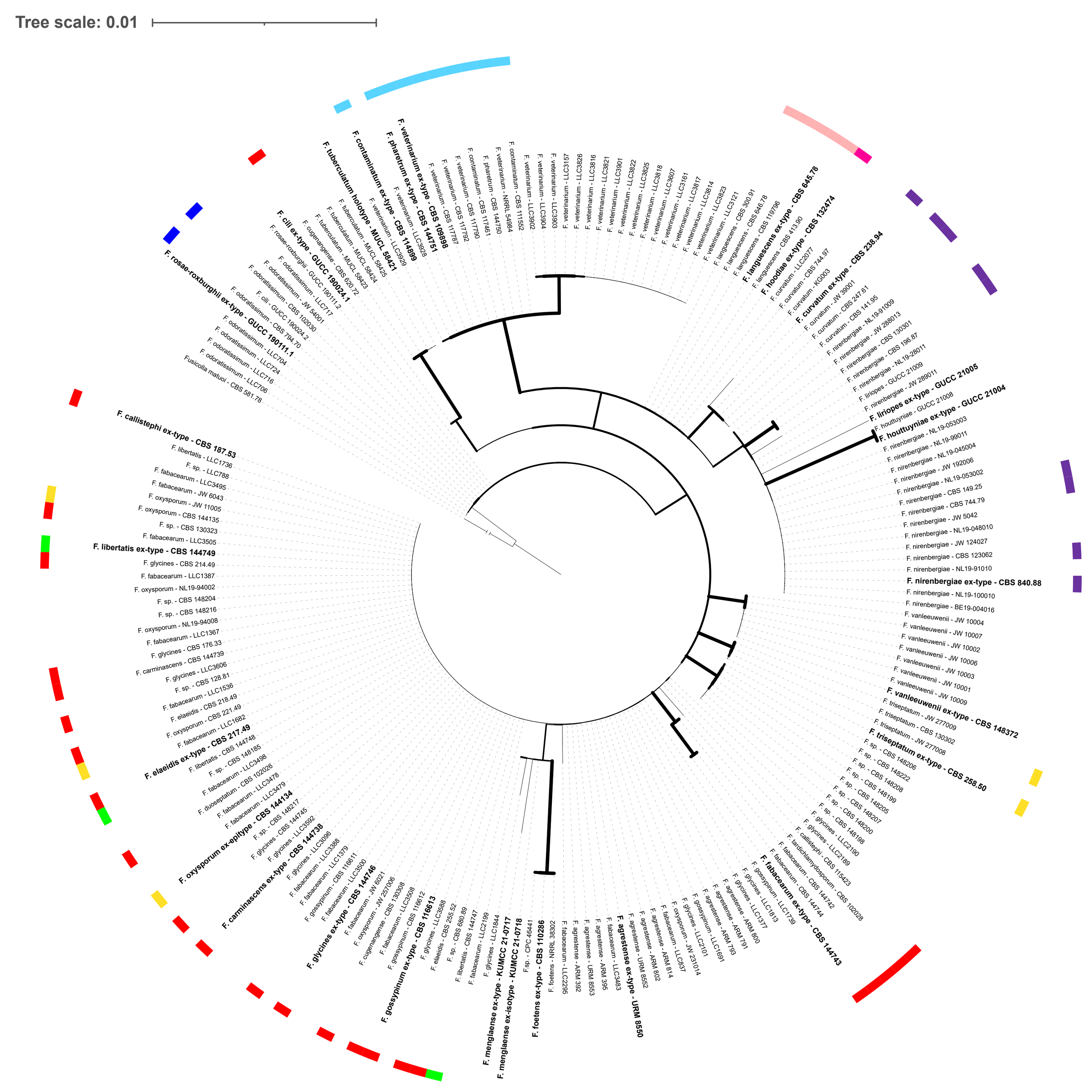


### Internal Transcribed Spacer (ITS)

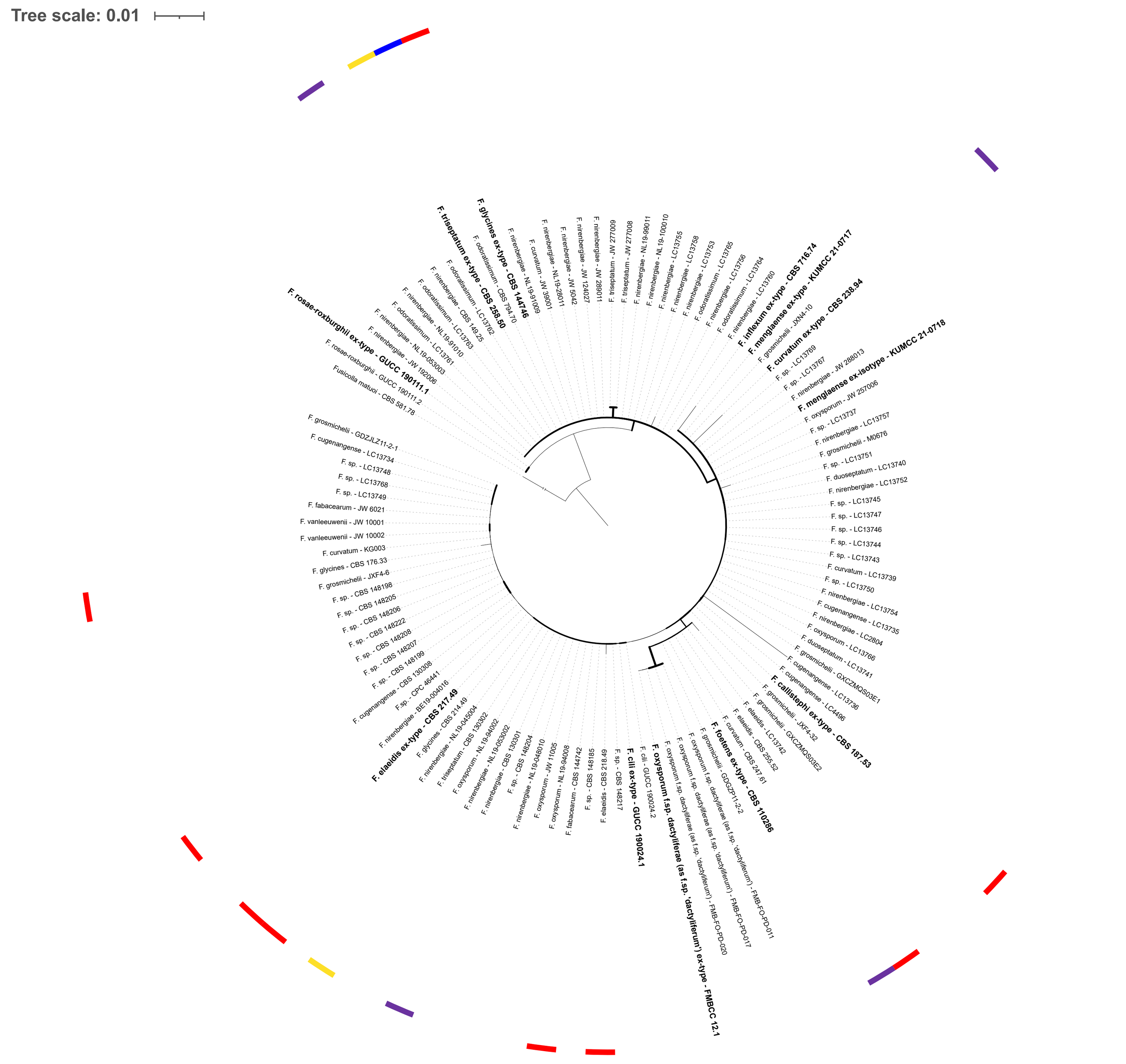


### RNA polymerase II largest subunit (*rpb1*)

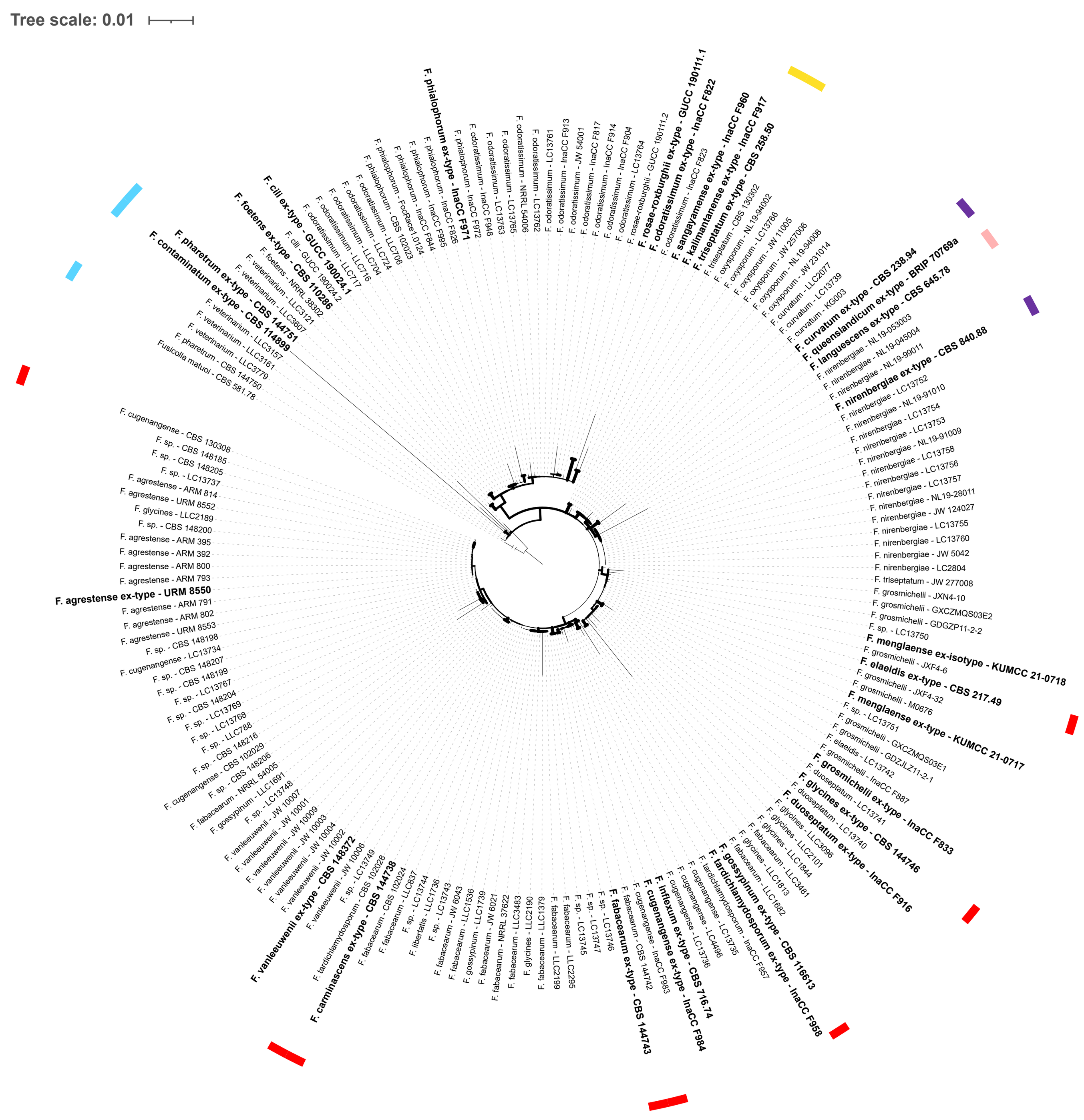


### RNA polymerase II second largest subunit (*rpb2*)

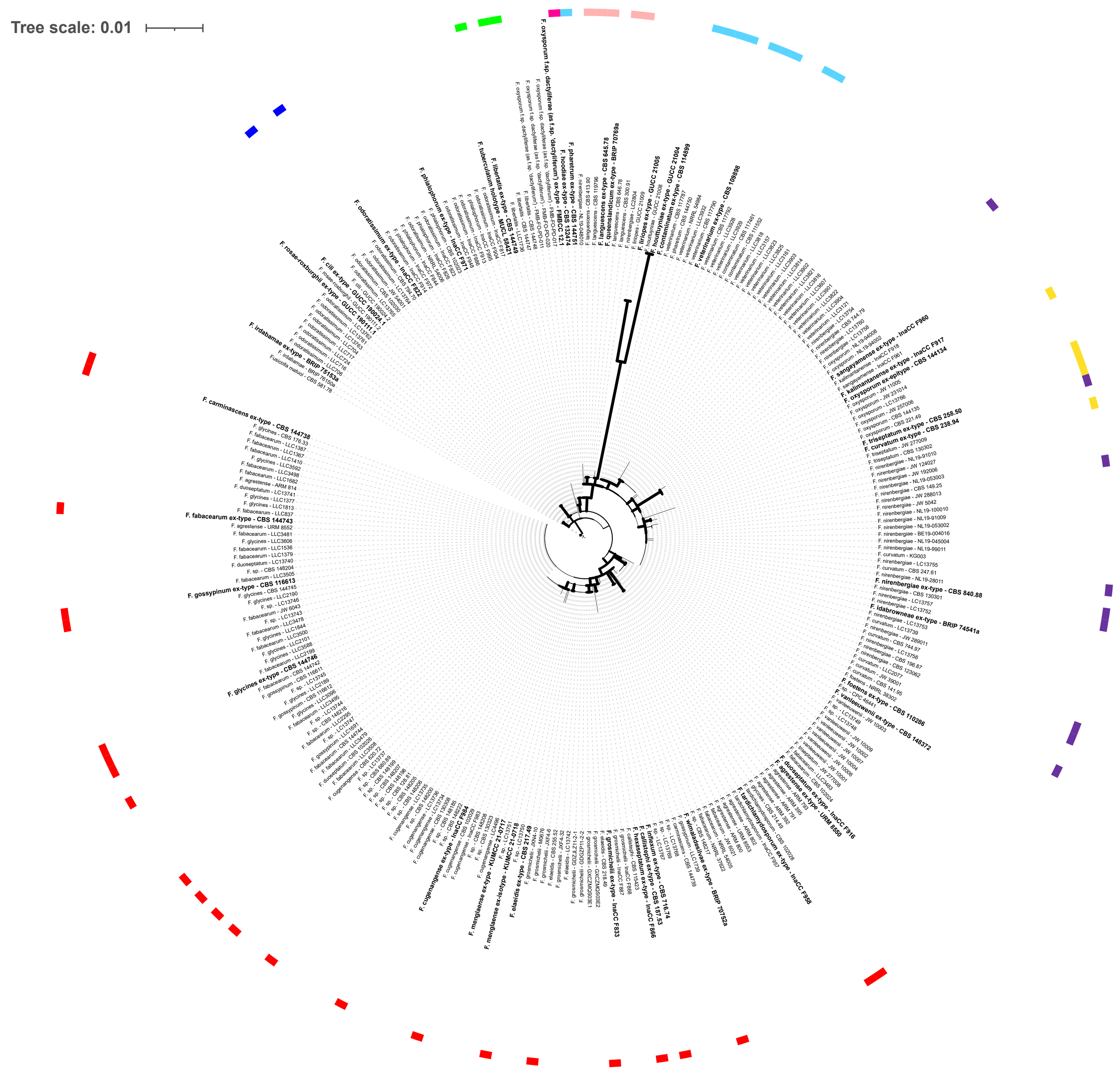


### Transcription elongation factor 1-alpha (*tef1* / *EF-1a*)

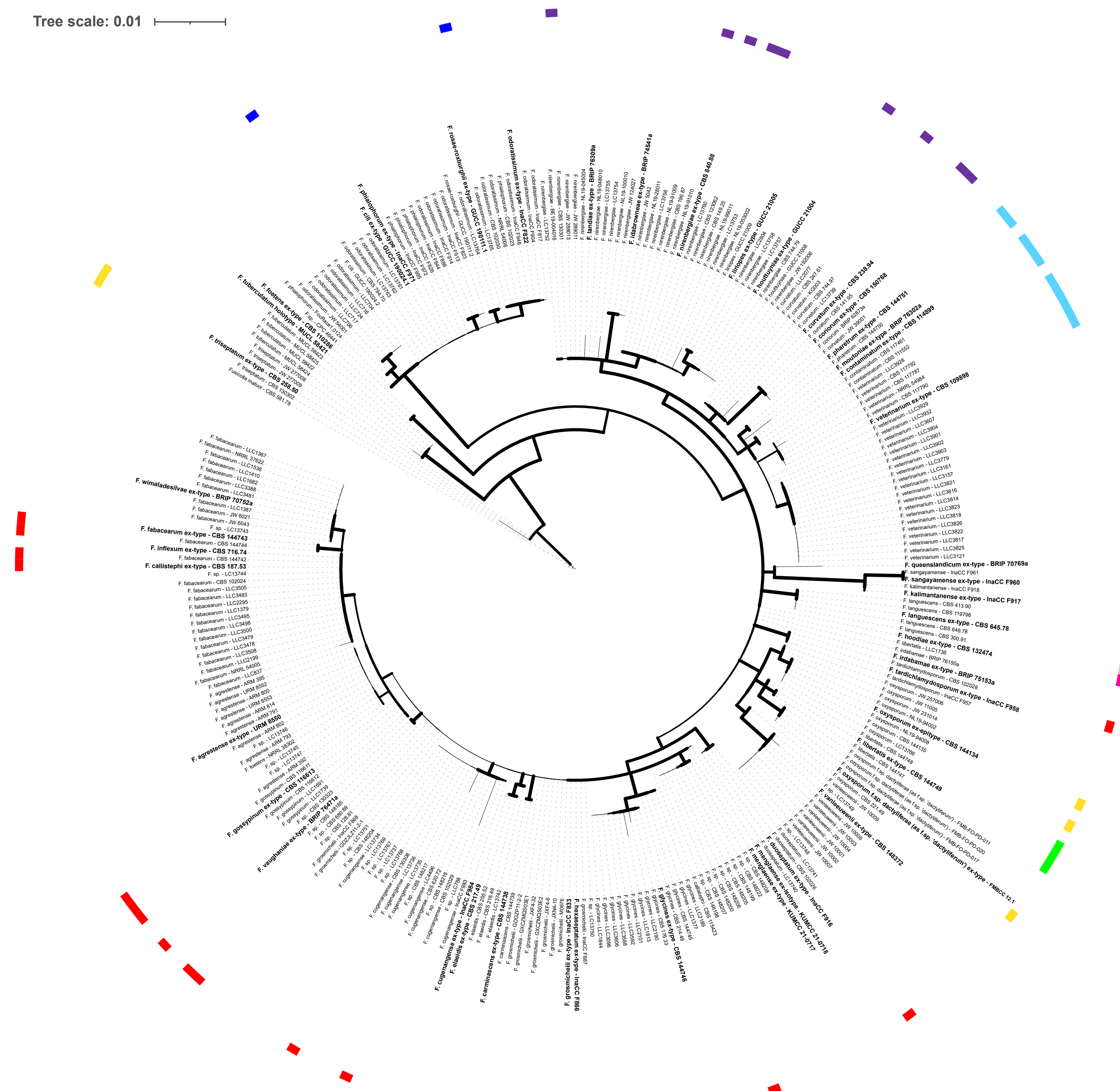


### Beta-tubulin (*tub2*)

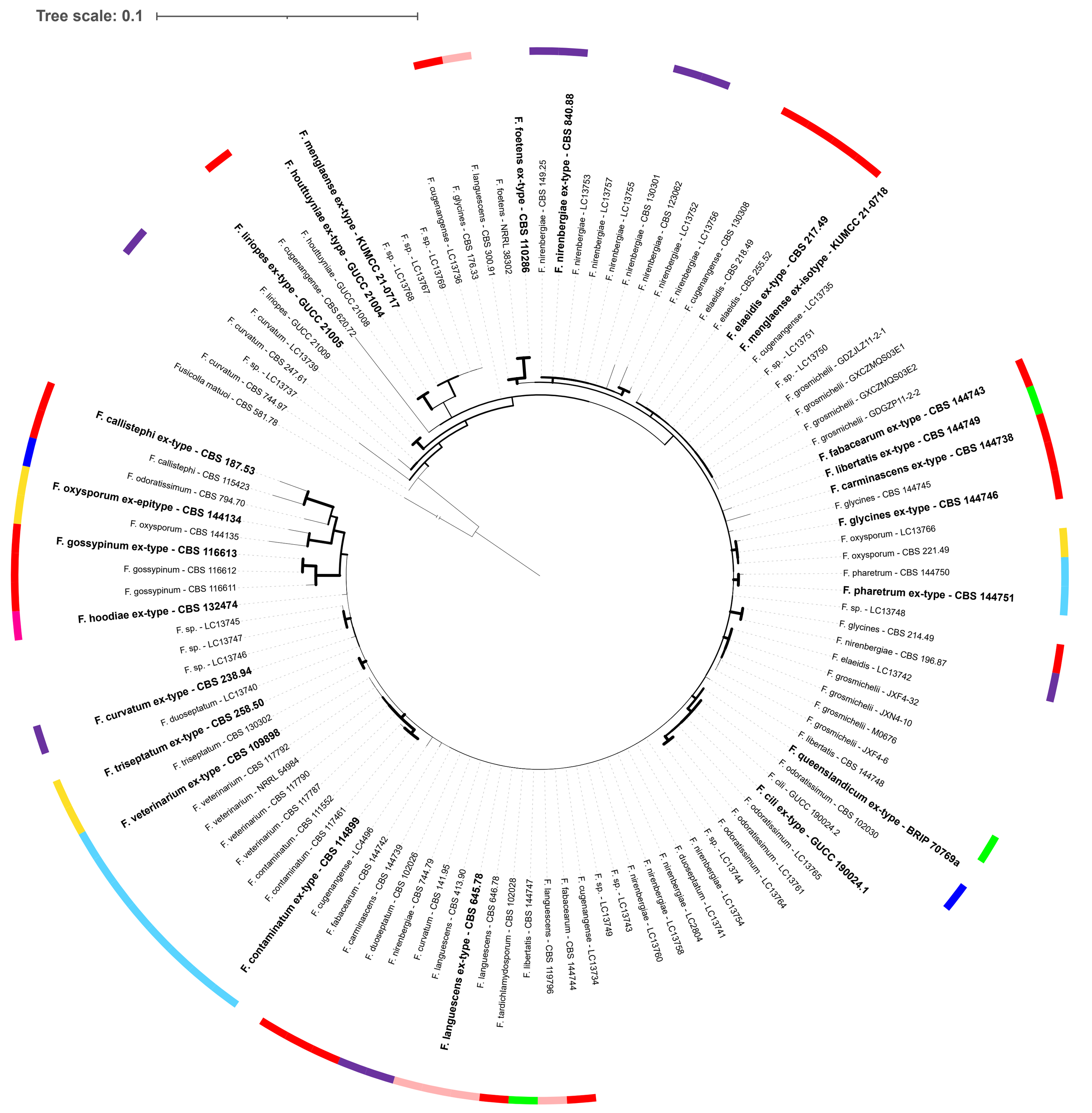


**Previously described clades**

| | Lombard et al., 2019 | O'Donnell et al., 1998 & 2004 | Laurance et al., 2014 | Brankovics et al., 2017 |
|---|---|---|---|---|
| (blue) | I | 1 | 1 | 1 |
| (green) | II | | | |
| (red) | III | 2 | | 2 |
| (magenta) | IV | | | |
| (yellow) | V | 4 | 4 & 5 | |
| (pink) | VI | | | |
| (light blue) | VII | | | |
| (purple) | VIII | 4 | 3 | 3 |
