## Supplementary figures and images for "Consolidating species boundaries in the *Fusarium oxysporum* species complex based on genomic evidence"

### Figure S3

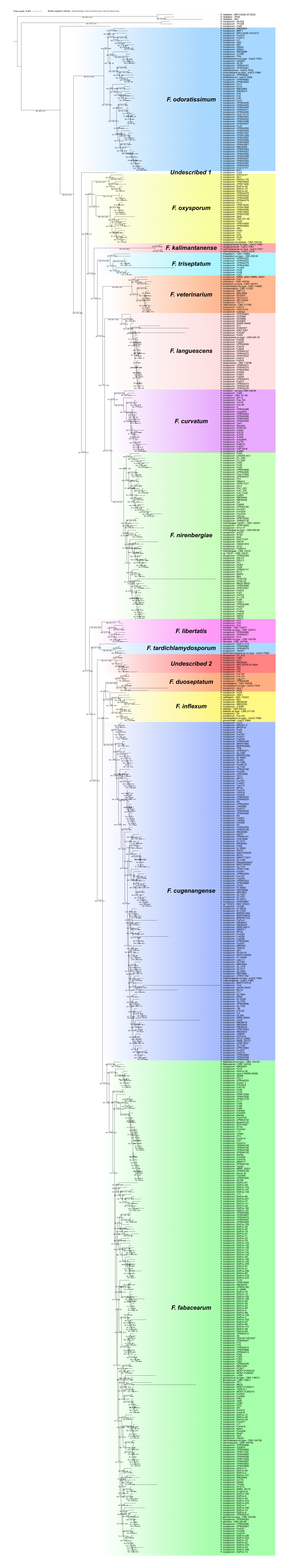

### Figure S4

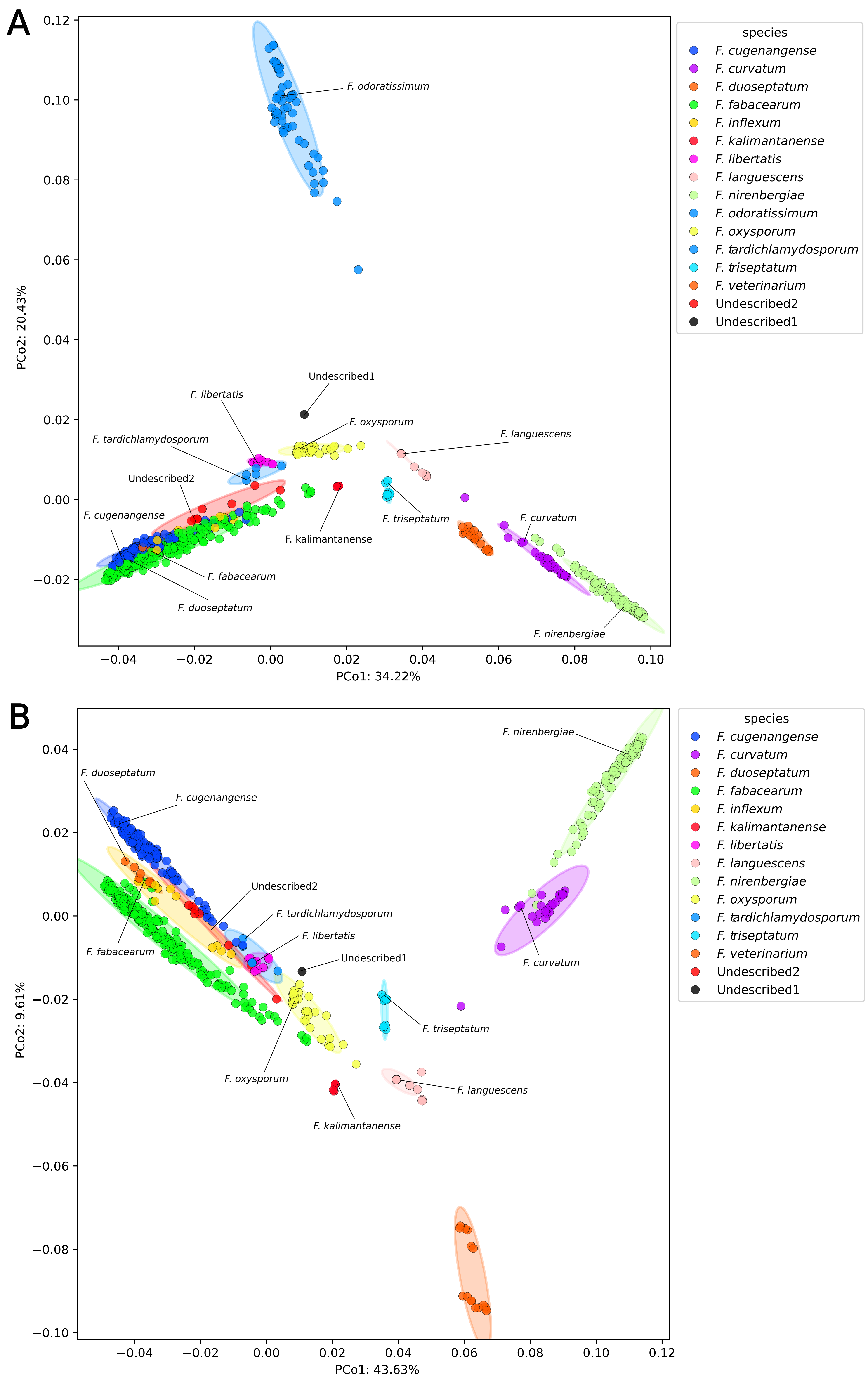
